# FGF and MafB regulated cadherin expression drives lamina formation in the auditory hindbrain

**DOI:** 10.1101/2025.04.30.651488

**Authors:** Rosanna C. G. Smith, Maryam Clark, Mireya Vazquez-Prada, Marc Astick, Kristina C. Tubby, Stephen R. Price

## Abstract

The avian auditory brainstem contains specialized nuclei critical for sound localization, including the nucleus laminaris (nL), which forms as a single-cell-thick lamina essential for computing interaural time differences. Despite its functional importance, the molecular mechanisms guiding nL lamina formation have remained poorly understood. Here, we identify a signalling cascade involving FGF8, MafB, and cadherin-22 that orchestrates this morphogenetic process.

We show that FGF8 is selectively expressed in the developing auditory hindbrain and correlates spatiotemporally with lamina formation in the nL. Disruption of FGF signalling—either via misexpression of FGF8 or dominant-negative FGFR1—perturbs the formation of the nL and alters cadherin-22 expression. In vitro culture experiments further reveal that nL lamination is sensitive to FGF8 dosage, with an optimal concentration required for both FGF8 and MafB expression and correct structural organization.

We demonstrate that FGF8 induces MafB, which in turn regulates cadherin-22 expression. Functional disruption of cadherins impairs lamina formation and leads to reduced FGF8 expression, indicating a feedback loop between adhesion and signalling. Cadherin protein expression appears enriched in the dendrites of nL neurons and computational models—both static and dynamic—show that bipolar, dendrite-localized, adhesion can drive laminar architecture as the maximum adhesion configuration.

These findings establish a novel molecular and biophysical mechanism for neuronal lamination in the vertebrate hindbrain, showing how local FGF signalling, transcriptional regulation, and dendritic adhesion converge to shape neural circuitry essential for sound localization.

## Introduction

A body of evidence suggests that the establishment of neuronal connectivity patterns during embryo development are, in part, a reflection of the settling position of neuronal cell bodies (Balaskas et al 2019, Hilde et al., 2016, Sürmeli et al., 2011, Tripodi et al., 2011). For example, monosynaptic proprioceptive sensory inputs to motor neurons have been shown to be dependent on the trajectory of sensory axon pathfinding and the position of motor neuron soma centrally. Dendritic morphologies have also been suggested to constrain the number, type, and distribution of pre-synaptic inputs on post-synaptic neuronal partners (Kostadinov and Sanes, 2015, Landgraf et al., 2003, Vlasits et al., 2016, Vrieseling and Arber, 2006). Moreover, positional features of axonal growth and segregation have been invoked as a strategy, independent of cellular and molecular recognition programs, to explain how pre-synaptic neurons find their appropriate targets (Langen et al., 2015). Together, these studies suggest that positional elements are critical in regulating the assembly and function of neuronal circuits. Therefore, during embryo development, it is crucial that neurons become positioned correctly and juxtaposed appropriately to facilitate appropriate synaptic contact during neuronal circuit formation.

A major mode of organization of neurons in more evolutionary ancient areas of the vertebrate central nervous systems clusters soma of functionally related neurons into groups known as neuronal nuclei. However, the mechanisms that drive the formation of neuronal nuclei, a process termed nucleogenesis (Agarwala and Ragsdale, 2002), remain largely enigmatic. The clustering of neuronal cell bodies into neuronal nuclei results in the formation of a variety of types of three-dimensional shapes, from ovoid and allantoid to layered structures. A particularly notable example of the different shapes that neuronal nuclei can form is found in the second and third-order nuclei involved in sound source localization in birds and reptiles (Boord, 1969; Book and Morest, 1990; Carr and Konishi, 1990; Joseph and Hyson, 1993; Knudsen and Konishi, 1979; Rubel and Parks, 1975; Rubel et al, 1976). These nuclei, the nucleus magnocellularis (nM) and nucleus laminaris (nL) are critical to the calculation of differences in sound arrival at each ear, the so-called interaural time difference (ITD), which is the major cue for low frequency sound localization in the azimuthal plane (Carr and Boudreau, 1993; Grothe et al, 2010; Hyson, 2005; Kuhn, 1977; Rayleigh, 1875; 1907). nM cell bodies cluster into an ovoid structure whereas nL cell bodies form a flat sheet of cells, in its mature formation, one single cell thick.

As summarized in Figure 1, the nucleus magnocellularis on each side of the brainstem receives synaptic input from the ipsilateral ear. nM neurons make axonal projections to neurons of both nuclei laminaris with ipsilateral synapses occurring on the dorsal surface of the nL sheet and contralateral synapses occurring on its ventral surface. The formation of the nL as a sheet of cells, in most places one cell-thick, is essential for coincidence detection of ipsilateral and contralateral synaptic inputs in allowing a calculation of the ITD for a given frequency of sound (Carr and Konishi, 1988; 1990; Jeffress, 1948; Smith and Rubel, 1979).

**Figure 1.**
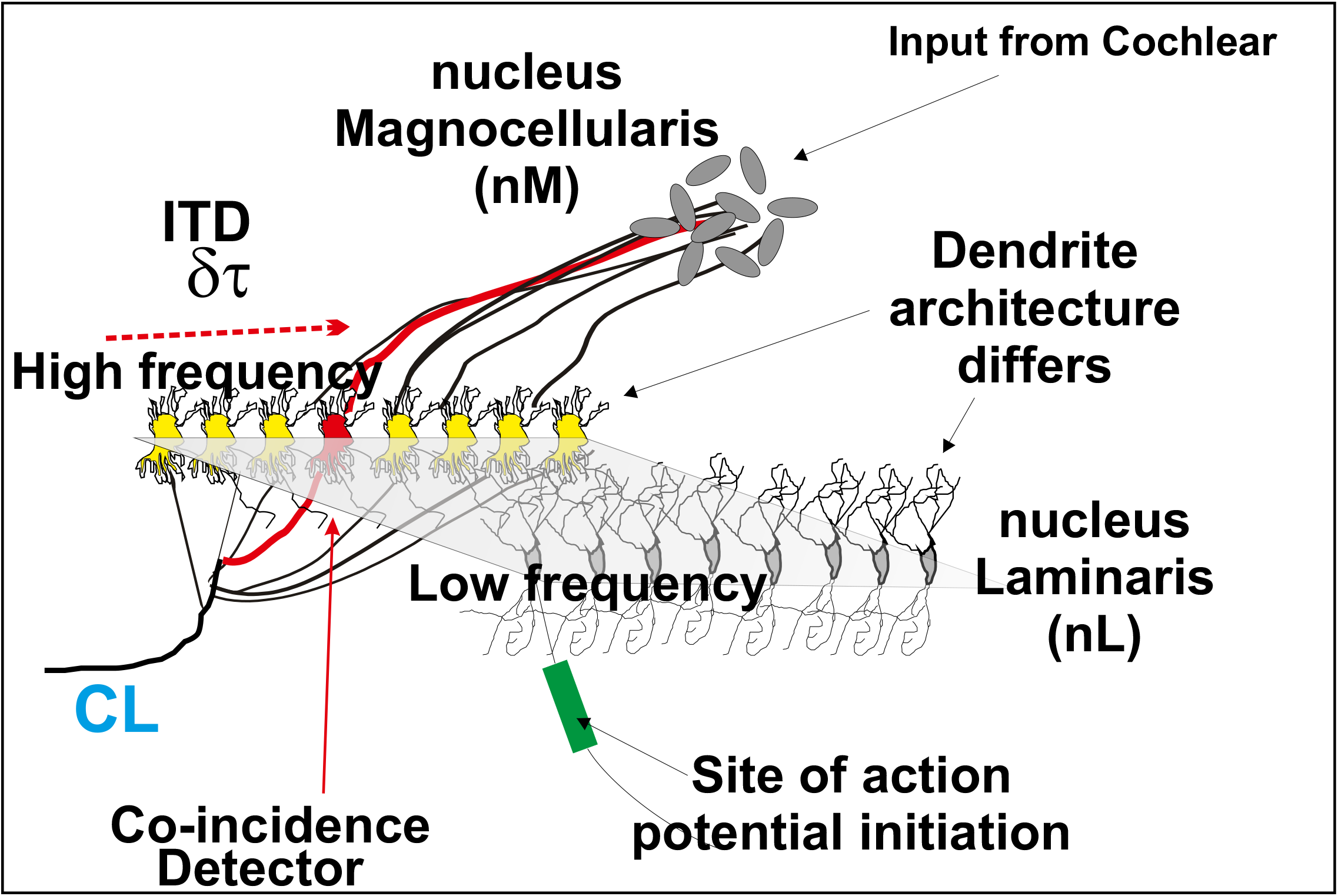
Schematic of the nucleus magnocellularis inputs to the nucleus laminaris (nL) and topographical organisaton of the nL. Bilateral nM inputs to dorsal and ventral dendrites of nL neurons create a neuronal map of auditory space. Coincidental ipsi– and contra-lateral inputs to a restricted subset of nL neurons, related to the interaural time difference (ITD) increase the probability of nL neuron firing an action potential (AP). The site of AP initiation and dendrite architecture of the nL neurons varies with characteristic frequency of the nL neuron.

The synaptic inputs to the nL are also patterned in the medio-lateral and the rostro-caudal extent of the flat sheet of the nL. Sounds of higher characteristic frequency are processed in the rostral-most extent of the nL. Further, synaptic inputs across the ventral dendrites of the nL in the medial-lateral extent of the nL represent sound source localization from closest to the midline to most laterally, respectively (Kubke and Carr, 2006). Thus, the structure of the nL as a lamina is essential to the integrity of sound source localisation with different frequencies being processed along the rostro-caudal axis and the azimuthal representation of sound source localization occurring in the medio-lateral extent of the lamina.

Despite the importance to the functioning of the avian auditory hindbrain of the physical shape of the nucleus laminaris as a flat sheet of cell, relatively little is known of how the morphology of the nL forms during embryo development. Part of the reason for this may be due to the lack of clear molecular markers during the timecourse of nL formation, before the nL can be identified from its lamina structure alone.

Previous studies of the morphology of the nL and nM show that both nuclei arise from the so-called auditory anlage within the dorsal hindbrain (Cramer et al, 2000; Tan and LeDouarin, 1991). The initial segregation of the nM from nL occurs early in auditory hindbrain development with the subsequent clustering of the nM soma and nL soma lamina formation coinciding with the elaboration and refinement of the dendrite architecture of both nM and nL cells (Hyson, 2005; Ohmori, 2014; Parks et al 1987). nM neurons initially elaborate radially symmetric dendrites which then degenerate to reveal a Calyx of Held configuration (Jahveri and Morest, 1982). In contrast, nL neurons elaborate a bi-tufted dendrite architecture that is initially uniform across the forming nL but refines to reveal short highly-branched dendrites more rostrally with longer less-branched dendrites more caudally (Smith and Rubel, 1979; Smith, 1981; Agmon-Snir et al, 1998).

In this study, we explore the molecular mechanisms responsible for nL formation and laminar organization. Specifically, we show that levels of Fibroblast Growth Factor (FGF) signalling result in expression of the transcription factor MafB within the nL (Eichmann et al, 1997). MafB expression subsequently induces expression of cadherin cell adhesion family members (Duguay et al, 2003; Hirano and Takeichi, 2012), which we show as being essential to the nL’s structure. We further suggest that cadherin function occurs within the dendrites of the nL cells and model how this could result in lamina formation.

## Materials and Methods

### Chick Embryo Preparation

Fertilised Brown Bovan Gold Hen’s eggs (Henry Stewart Farms, UK) were incubated in a humid, forced draft incubator (Lyons Inc) at 39°C and staged as in Hamburger and Hamilton (1992; 1951) (HH)). All embryos were treated in accordance with the Animals (Scientific Procedures) Act of 1986, UK and its amendments.

### DNA Constructs

The plasmid for generating a cRNA probe to chicken FGF8 was a kind gift of H. Ohuchi, University of Tohushima, Japan. Constructs for cRNA probes to cadherin-2, cadheri-13 and cadherin-24 were generated by PCR from cDNA prepared from e2 to e4 chick embryo mRNA. Antisense digoxygenin (DIG) RNA probes were generated using standard techniques Full-length cDNAs for chick cadherin-2, GFP or NΔ390 were cloned into a pCAGGS vector. Other constructs used in this work include transposase integrated doxycycline inducible N-cadherinΔ390 plasmids (Tanabe et al, 2006). *MafB* and *dnMafB* vectors (Suda et al 2014) were a kind gift from M.Tanaka (Tokyo Institute of Technology), mFGF8 electroporation plasmid was the kind gift of T. Shimogori (Fukuchi-Shimogori and Grove, 2001) and the dominant negative FGFR1 construct (Saffell et al, 1997) was a kind gift of I. Mason (KCL).

### In Situ Hybridization Histochemistry

Digoxigenin (DIG)-labelled anti-sense cRNA probes were used for in situ hybridization histochemistry on 15 μm thick cryostat sections as in Price et al 2002. Dual in situ hybridization with immunohistochemistry was performed by replacing the proteinase K treatment with a 30-minute incubation of the slide sections with 0.1% Triton X-100 in phosphate buffered saline (PBS) (pH7.4). Development of the RNA in situ was as normal followed by incubation of the slides with 4M HCl for 5 minutes with subsequent washing for 15 minutes in PBS. The primary antibody for BrdU was then applied for 16 hours at 4°C. Following this, secondary HRP conjugated goat anti mouse antibodies were applied and ImPact DAB substrate reagent (Vector labs) was used to reveal HRP immunohistochemistry following manufacturers instructions.

### Bromodeoxyuridine labelling

Bromodeoxyuridine (BrdU) at 500μM or 125μM dissolved in Hanks balanced salt solution was applied via syringe application on top of staged and windowed embryos varying at start points between stages 18 and 27. Either a single dose was administered or two doses, 12 hours apart.

### In Ovo Electroporation

Expression of cDNAs was achieved by *in ovo* electroporation using an ECM830 electro-squareporator (BTX Inc.). ∼0.1μl of DNA construct (1-10μg/μl in H_2_O with 0.1% Fast Green (Sigma)) was pressure injected into the lumen of the brain stem. Five 30 Volt electrical pulses of 50ms duration equally spaced over a 5 second period were applied by placing electrodes adjacent to each side of the head of the embryo. Embryos were electroporated at HH stages 12-20 and analyzed at HH stages 29-32. For NΔ390 experiments, doxycycline hyclate (200μl) at a concentration of 0.25 μg/ml was applied from stage 20 onwards.

### Immunohistochemistry

Antibodies used in this study were: Rabbit (R) anti-GFP (Invitrogen, 1/1000), R anti Caspase 3 (cleaved) (Cell Signalling Technology), R anti-*N*-cadherin (1/1000; AbCAM) M (Mouse) anti-γ-catenin (1/100; BD Biosciences), M anti-GFP (1/100; Invitrogen), M anti-BrdU (1/50; Roche), 2D6 and 4D5 (M anti-Islet-1 from the Developmental Studies Hybridoma bank), Alkaline phosphatase conjugated Sheep anti-DIG Fab fragments (Roche) (1/5000). Immunocytochemistry was performed essentially as described in Price et al 2002. Cryostat sections mounted on superfrost plus glass slides were incubated in PBS for 5 minutes followed by incubation in block solution (PBS with 1% Goat serum (Sigma)) for 30 minutes at 20°C. This solution was replaced by antibody diluted in block solution and incubated for 12 to 16 hours at 4°C. Following three washes of 5 minutes in PBS, fluorescent conjugated secondary antibodies were incubated with the sections for 30 minutes at 20°C in block solution, washed and mounted with vectashield fluorescent mounting medium (Vector Labs).

### Brainstem Slice Cultures

Slices were cultured following a modified protocol of Sanchez et al, 2011. The chick embryo is removed from the egg at the appropriate stage and dissected for the hindbrain in Pannett Compton solution on ice. Dissected hindbrains were then placed in 4% low melting point agarose (Thermofisher) and fixed via superglue to a Leica VT100S vibratome stage, anterior side down. Glued agarose is then placed in 5% CO2 equilibrated chicken artificial cerebrospinal fluid (ACSF) on ice. 600μm vibratome slices were dissected from the agarose gel using clean forceps and placed in a medium sized petri dish on ice in equilibrated ACSF. Slices are taken to electroporation table consisting of a dissecting microscope and an electrosquare porator (BTX ECM830). Plasmid was placed on the slice and electrodes are then placed either side of the slice and current is activated. Electroporated slices are immediately put in chicken 1-2ml of culture media fortified with varying concentrations of FGF8 (Human-murine FGF8, CN:100-25, Peprotech) and placed in sterile 5% CO2 incubator and cultured at 37°C. Culture media is replaced every 10 hours with fresh media and plates are changed to avoid contamination of primary cultures.

### Image Acquisition and Analysis

Images were acquired on a Nikon Eclipse E80i fluorescence microscope equipped with a Nikon DS5M and Hamammatsu ORCA ER digital camera or on a Leica SPE confocal microscope. IPLab software (Scanalytics, Rockville, US) was used for merging false-colour images in multichannel fluorescent staining and ImageJ (National Institutes of Health, US, public domain) (Schneider et al, 2012) was used to make image measurements. Any image manipulation (for example contrast enhancement) was applied equally across all images under comparison. Lengths and areas as measured using ImageJ were calibrated using measurements of an object of known dimensions taken under the same magnification. Cell counts were made for nuclear BrdU-labelled neurons and the cytoplasmic caspase-3 or FGF8 stain. The Abercrombie (1946) correction was used for cell counts across more than one tissue section. For individual developmental stages, as our results are confirmatory of those published previously, we quantitated all neurons in one embryo at each stage. All other results are indicative of at least three embryos unless explicitly stated in the text.

### Automated nL Morphology Analysis

Initial image segmentation was carried out using ImageJ. Automation was achieved using the Python version 3.8.1, using the scikit-image 0.16.2, scipy 1.3.0, numpy 1.17.2, matplotlib 3.0.2 and pandas 1.0.5 packages. Briefly, the r^2^ number calculated for each section was calculated as a classic coefficient of determination approach. A total sum of squares (TSS) for the initial spread of cells was calculated from the central cell in the nL. Then a best-fit line was drawn through the nL and the residual sum of squares (RSS) of distances of each nL cell calculated from this best-fit, r^2^ was calculated as 1-RSS/TSS. An r^2^ close to 1 indicates an nL closely aligned to the best fit line, one close to 0 indicates an nL with larger variance away from the best fit (more like a ball of cells than a line of cells).

### Maximum Adhesion Model

Wolfram Mathematica 7 (Wolfram Research Inc, US) was used in graphing. Full details of the model are provided in Appendix 1 and 2.

### Dynamic Bipolar Model

The dynamic bipolar model (figure S7) was developed on Python version 3.5.1, using the numpy 1.17.2, matplotlib 3.1.1, scipy 1.4.1 and pytest 5.4 packages. We developed a simulation to visualize nL cell dynamics over time, incorporating mathematical modelling of their intercellular interactions and dendritic extensions.

Each cell was represented as a central body with dendrite-like focal points of two types: Type A (NfpA, red) and Type B (NfpB, green). These dendritic extensions were modelled as discrete points attached to the body, interacting both with the cell and with dendrites of neighboring cells.

The simulation updates cell positions iteratively based on a set of interaction forces. Specifically, the total force acting on each cell *i* at time *t* was computed as:

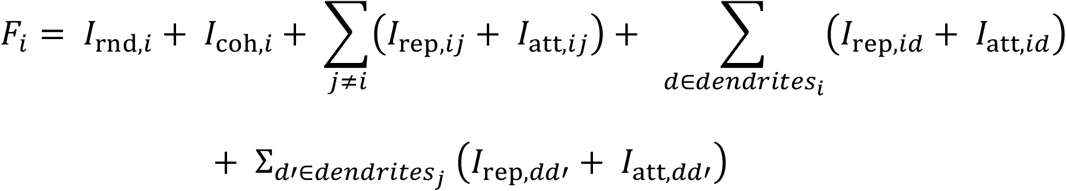

Where:

1. *I*_*rnd*_ represents random forces acting on cell *i*, introducing stochastic movement to the system
2. *I*_*coh*_ represents a cohesion force that biases cell movement towards maintaining proximity to nearby cells.
3. *I*_*rep, id*_ and *I*_*att*, ij_ describe repulsion and attraction between two different cell bodies, *i* and *j*.
4. *I*_*rep, dd’*_ and *I*_*rep, dd’*_ describe interactions between dendrites of different calls.
5. *I*_*rep, dj*_ and *I*_*att, dj*_ represents forces between a dendrite of one cell and the body of another cell.

The repulsion forces *I*_*rep*_ dominate at short distances to prevent overlap while attraction forces *I*_*att*_ act within an intermediate range, encouraging the formation of structured interactions. The forces decay beyond a maximum interaction distance *r*_*maxrange*_. The random force *I*_*rnd, I*_ introduces variability in motion, and the cohesion force, *I*_*coh, I*_ ensures that cells do not disperse indefinitely.

At each time step *Δt*, cell positions *r*_*i*_*(t)* were updated according to:

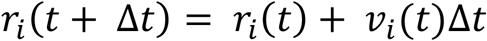

Where velocity *v*_*i*_*(t)* evolved based on the net force:

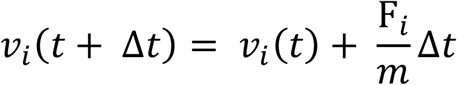

Assuming unit of mass (m = 1) for simplicity.

The simulation captured nL cell movement over time, where cells were represented as blue circles and dendrites as smaller circles (red for Type A and green for Type B) connected by arrows.

## Results

### Developmental profile of MafB expression in the auditory hindbrain

To investigate formation of the nucleus laminaris we first sought a molecular marker of the auditory hindbrain that would allow us to follow its formation over time. Previous studies have identified the expression of the basic leucine zipper protein MafB in the mature avian auditory hindbrain, including the second order auditory nM and nucleus Angularis (nA) as well as the third-order nucleus Laminaris (nL), amongst other neurons in the brainstem (Eichmann et al, 1997). The reported expression of *MafB* (was at E12 (HH Stage 38) in the developing chick, after the lamina structure of nL had already forms. MafB has also been shown to have a prominent expression profile in rhombomre 5 and 6 from HH st ∼9. We therefore investigated the developmental profile of *MafB* in the auditory hindbrain by antisense RNA in situ hybridization from HH stage 26 onwards, around when the auditory anlage is first identifiable (Figure 2 A-F) (Book and Morest, 1990). At HH stage 26, *MafB* was expressed in the presumptive nucleus Angularis of the auditory hindbrain along with a prominent expression in cells more medial to the nA (Figure 2A). This pattern of expression persisted to HH stage 30 (Figure 2 B, C), with the *MafB* expression in the nA in increasing numbers of cells. However, at st 30, we observed expression of *MafB* in an ovoid cluster of cells more dorsal to the nA, which we define as the presumptive nL. This pattern of expression of *MafB* in the nA and nL was confirmed at later stages (HH st 34, st 35 and st 39; Figure 2D-F) but we also began to detect expression of *MafB* in the presumptive nM from HH st 35. The expression of *MafB* in the nL mirrored the lamina structure of the nL from HH st 34 onwards with a final, flattened sheet observable at HH st 39. We thus confirmed expression of *MafB* in the nA and nL and a lower level of expression in the nM. However, we considered that the prominent expression of *MafB* in the nA cells in addition to nL from stage 30 and nM from st 35 might be problematic as a marker for studying nL formation as a lamina.

**Figure 2.**
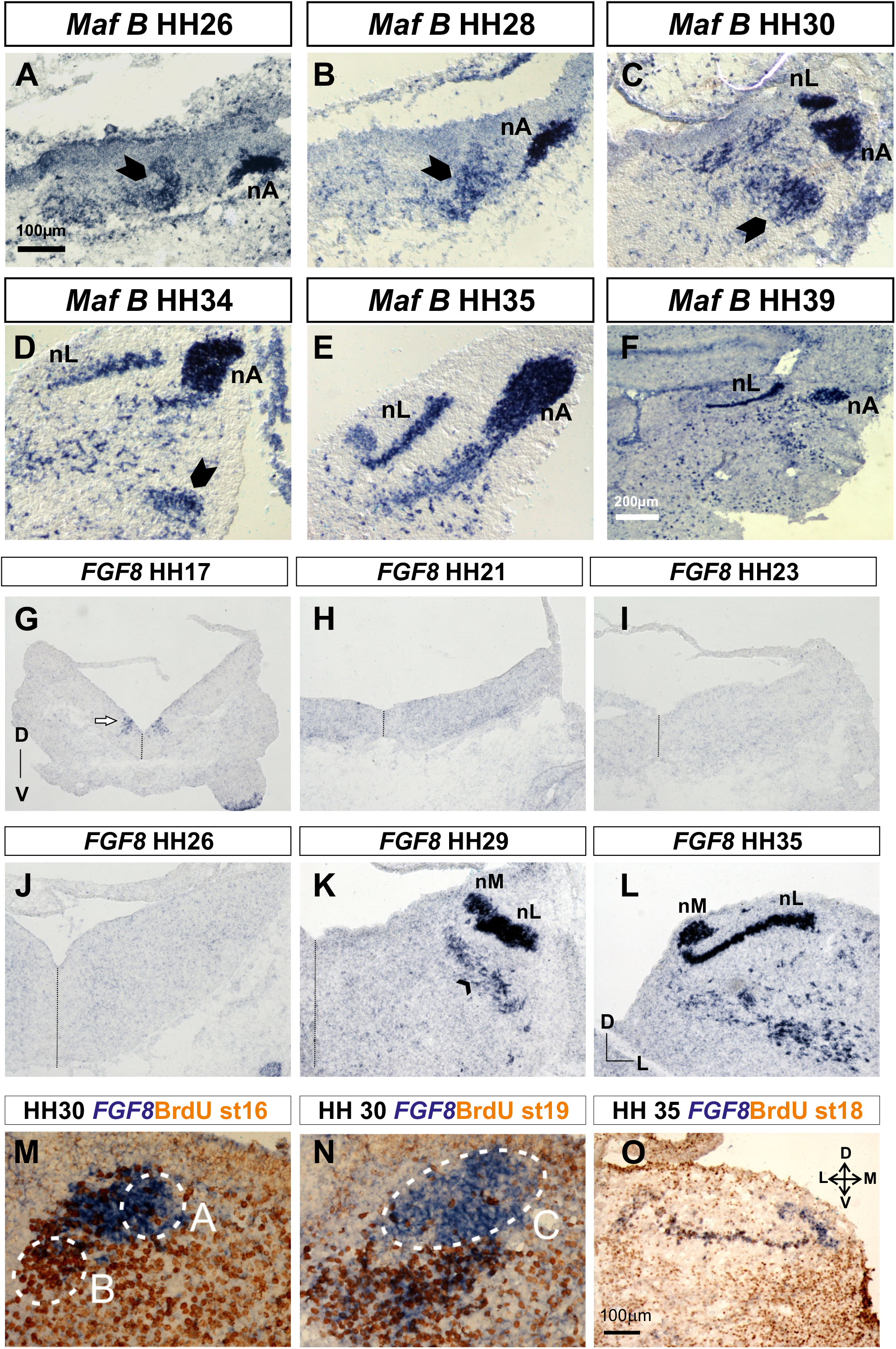
A-F, *mafB* expression at various HH stages indicated in the boxes above the panel. G-L FGF8 expression at the HH stages indicated in the boxes. M, N *FGF8* and BrdU expression at HHst30 following BrdU application at st 16 (M) or st 19 (N) O FGF8 and BrdU expression at HH st 35 following BrdU application at HH stage 18.

We thus decided to take a candidate molecule approach to search for a molecular marker for nucleus laminaris formation that was more regionally restricted in expression. Previous studies have identified FGF signalling as being important in the expression of *MafB* in the early stage otic/pre-otic regions of the brainstem (Marin and Charnay, 2000). We thus investigated FGF family expression in the auditory hindbrain from the earliest stages of nM and nL generation (from HH stage 17 (Figure 2G-O)).

### *FGF8* is expressed in the nucleus magnocellularis and nucleus laminaris and follows lamina formation

During our investigations, we found that FGF8 is expressed within the presumptive developing auditory anlage of dorso-lateral rhombomere 5 from Hamburger Hamilton (HH) stage (st) 28 and expression persisted nM and the nL until at least HH st40, the latest developmental stage studied (Figure 2 G-O, Fig S1-S5). The nL and nM can be identified by their respective laminar and clustered structures once they have developed, but the idenfication of a molecular marker for consituent cells of the late auditory anlage and presumptive nL and nM allowed us to characterize the morphological and molecular development of the auditory localization circuitry. Prior to its expression in the auditory anlage at stage 28, FGF8 expression was observed in a small number of scattered cells in the dorso-medial hindbrain from HH st 26-29 (Figure 2J, K) and by stage 30, the auditory anlage was clearly distinguishable by FGF8 expression (Figure 2M-O; Fig S2). The auditory anlage, as delineated by FGF8 expression at this stage, appeared roughly allantoid in shape over three dimensions. By HH st33 (Fig S3), the separation of nM and nL assessed by FGF8 expression was clear from ∼160μm from the rostral most extent of FGF8 expression. By this stage, the shape of the nL shape, as assessed by FGF8 expression appeared to be flattening rostrally. This flattening of the nL was most apparent at rostral HH st36 (Figure 2L; Fig S4), where the nL was around 3 cell diameters thick and roughly lamina in shape

The total rostrocaudal extent of FGF8 expression in the brainstem was ∼600μm at both HH st33 and HH st36. By stage 39 (FigS5), lamina formation in the rostral nL is complete with a single-cell thick layer forming in the rostral most ∼300μm. The remaining caudal nL appears to continue flattening into a lamina structure after stage 40 (Data not shown). Cell counts of presumptive nM and nL during these stages indicated a steady increase in number of FGF8 positive cells from stage 30 to st36 for both nL and nM (∼1500 nL, ∼2000nM st30 to ∼4500nL,

∼6300nM at st36). Consistent with previous studies, we observed a decrease in both nM and nL between st36 to st39, the period at which apoptosis has been reported in both nuclei (∼3400 nL and ∼5700 nM at st 39). However, in the regions of nL rostral to nM between st30 to st36, a relatively constant number of neurons was found (880 cells st 30 to 812 cells st36). This rostral region of the nL is where a wave of lamina formation occurs between st 30 to st36, suggesting that lamina formation is independent of neuron number.

### nL and nM cells are segregated in the early auditory anlage

We next sought to address the positioning of nM and nL cells within the auditory anlage prior to separation of the two nuclei. Previous fate mapping studies have shown that these nuclei of the auditory brainstem have different but overlapping birth dates of their constituent neurons (Cramer et al, 2000; Rubel et al, 1976).

Neurons of the nM are born before around st17 with the majority of nL cells being generated between st17 and st22. We thus applied BrdU to embryos from HH st 16 through to HH st27 and assayed nM and nL formation through expression of FGF8 at HH st 29 and HH st 35 (Figure 2M-O; Fig S1). BrdU application to embryos at st16 followed by FGF8 RNA identification at st29 shows that the presumptive nM lies at the dorsomedial extent of the auditory anlage (Figure 2M). BrdU application at stage 19 confirmed this identification with the majority of staining in the auditory anlage being ventrolateral indicating the presence of the nL as a coalesced group of cells (Figure 2N). This suggests that within the auditory anlage, nM cells and nL cells already occupy distinct positions and little intermingling of the cells occurs. We also confirmed a rostral to caudal generation of nL cells through application of BrdU at stages between st18 and st22 followed by identification of FGF8 expression at st35 (Fig S1). Together, these results suggest that FGF8 represents a clear marker for nL and nM segregation within the auditory anlage and that nL cells form an ovoid structure prior to flattening to a lamina between st30 and st36. We thus sought to analyse the formation of the nL to observe segregation in more quantitative detail.

### Quantitation of lamina formation of the nucleus laminaris

To attempt to quantitate lamina formation, we sought a measure of the shape changes of the FGF8 expression within the nucleus laminaris from stage 30 to stage 36 (Figure 3A-G). Lamina formation was quantitated by measuring the length (medial to lateral extent) and width (dorsal to ventral extent) at several points on each of every fourth section of the nL at each of st 30, 33 and 36, as described in the schematic shown in figure 3H and the stage 33 examples in figure 3E-G. This approach was taken as nL lamination was observed to vary both rostrocaudally and mediolaterally.

**Figure 3.**
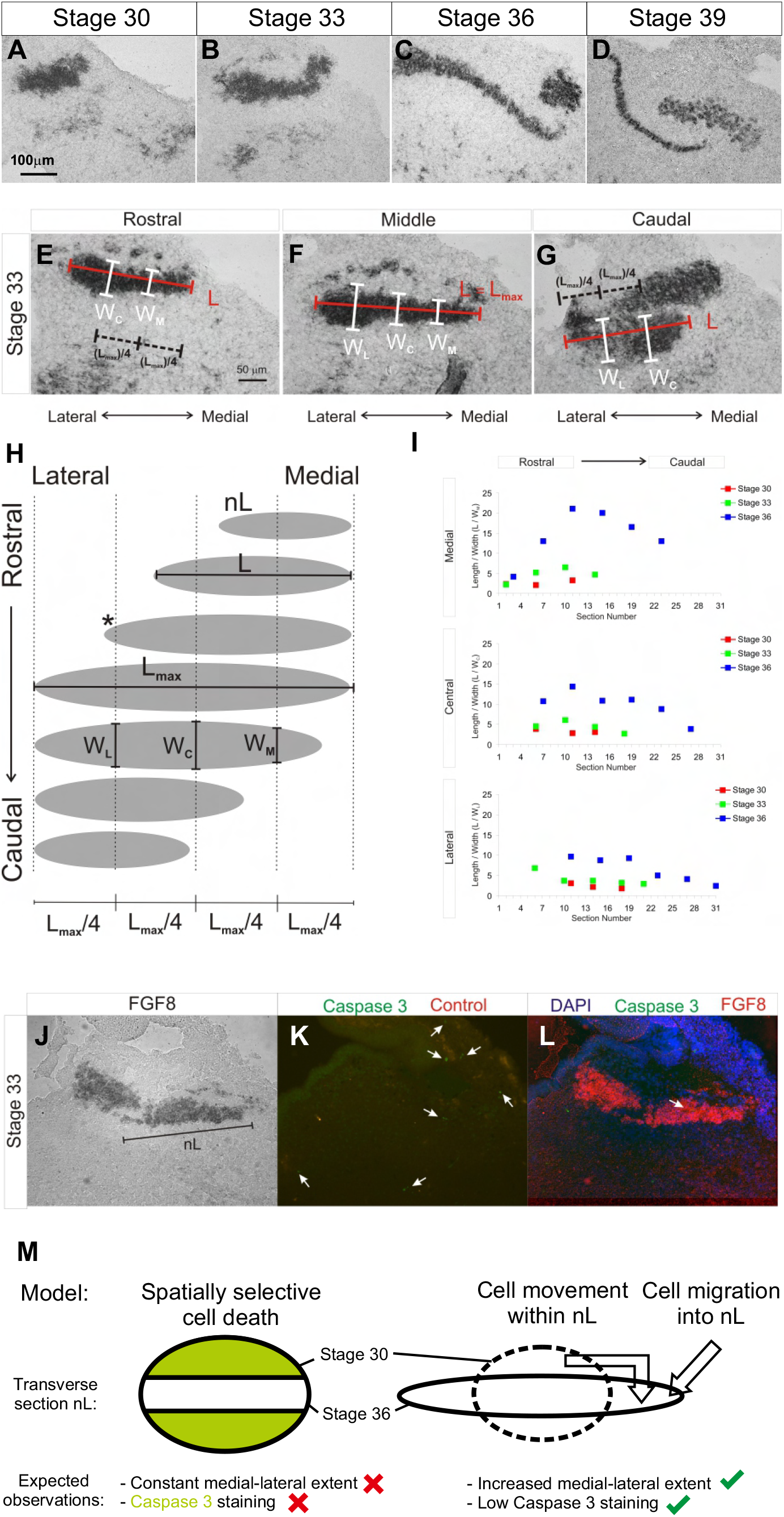
A-D FGF8 expression at HH stage 30-39 indicated in the boxes above the panels. E-G FGF8 expression at HH stage 33 annotated to indicate the medial (Wm) Central (Wc) and Lateral (Wl) widths of the nL and the Lengths (L) of the nucleus. H schematic of that shown in E-G. I Graphs showing length to width ratios of the nL in Medial, Central and Lateral regions at HH stages 30, 33 and 36. J-L FGF8 expression at HH st 33 (J,L) related to Caspase-3 expression in the nL compared to a control area of the hindbrain (K) M potential models of lamina formation of the nL by spatially restricted cell death or cell movement.

Stage 30 measurements do not show much variation in length/width values either rostrocaudally, for a given measurement position, or across the measurement positions. At HH st 30, the nL nucleus is approximately three times as long as it is wide (Figure 3I). Stage 33 shows more variation in length/width with slightly higher values than at stage 30 for medial and central measurements but similar values for lateral measurements. The greatest change in structure can be seen by comparing the stage 30 and 33 measurements with stage 36 (Figure 3I). For stage 36, the variation is greatest, with medial length/width values being on average higher than central values, which in turn, are higher than lateral values. A decrease in length/width can also be seen for the rostral to caudal axis although all length/width values from the medial position are greater than any in the lateral position, confirming the rostromedial to caudolateral lamination of nL during this period of development.

Thus, quantitation of lamina formation in these rostral regions through measuring the ratio of length to width of nL in medial, central and lateral regions of the nL suggested that the flattening of the nL occurs in a rostromedial to caudolateral wave: From st30 to st36, particularly from st 33 to st36, the rostral nL progressively lengthens as its width decreases (Figure 3I).

Together with our cell-count measurements of FGF8 expression in the nL, these length and width ratios suggest that the nL lamina formation occurs via a movement of cells within the nL, rather than a selective removal of nL cells via cell death away from a lamina configuration.

However, we sought to confirm this by assessing a potential role for patterned cell death in the lamina formation of the nL. We assayed cell apoptosis in the auditory anlage via detection of activated caspase 3 immunofluorescence with concurrent FGF8 expression via fluorescence in situ hybridization (Figure 3J-L). In this experiment, we detected a low incidence of activated caspase 3 equivalent to less than 0.45% of cell death at any stage between st30 to st36. Taken together, these results suggest that cell death is unlikely to explain the flattening of the nL from st30 to st36 and that spatial reorganisation drives the nL into a single-cell-thick sheet (Figure 3M). This spatial reorganization of the nL begins at HH st30, is ongoing at HH st33 and becomes mature as a lamina from HH st36 to HH st39. Further, a rostral to caudal and medial to lateral wave of lamination is observed.

### Cadherin/ catenin expression within the nucleus laminaris

The observed flattening of the nL from an ovoid structure into a lamina prompted us to ask what mechanism might drive such a shape change. Members of the cadherin family of cell adhesion molecules have been shown to drive coalescence of neuronal nuclei in the spinal cord and brainstem (Astick et al, 2014; Knufer et al, 2020). We thus first asked whether members of the type I and type II cadherins were expressed within the auditory hindbrain. We undertook a systematic assessment of the expression of five type I cadherins (cadherins 1-4 and the atypical type I cadherin-13 AKA T-cadherin) and ten type II cadherins (cadherin –5, 6, 7, 8, 9, 10, 11, 12, 20 and 22 (Hirano and Takeichi, 2012)) within the auditory hindbrain at various stages from HH st17 through to HH st39 (Figure 4A-O). Of these cadherins, we found that the type I cadherin cadherin-2 (Figure 4 J-Q) the type II cadherin cadherin-22 (Figure 4B, E, H) and the atypical type I cadherin-13 (Figure 4C, F, I) were expressed in the auditory brainstem. Cadherin-2 was found in both the nM and nL (Figure 4M-O; Figure 4Q) whereas cadherin-22 and cadherin-13 appeared expressed on only the nL and nA and not nM (Figure 4B-I). Both cadherin-13 and cadherin-22 expression in the auditory hindbrain was first detected at HH st30, around the time of nL expression of MafB and after the initiation of nM and nL expression of FGF8. This expression of cadherin-13 and cadherin-22 persisted until at least HH stage 39, the latest stage analysed (Figure 4B, C). The expression of cadherin-22 in the nL tracked the lamina formation similarly to FGF8 expression whereas we observed a reproducible difference in early expression of cadherin-13 which appeared graded across the nL with highest expression in more lateral regions of the nL (Arrows in Figure 4F, I compare to Figure 4D, E, G, H)). However, we detected cadherin-2 expression in a dorso-lateral cluster of cells around rhombomere 5/6 from HH st21 (Figure 4K-M), just after the generation of the nM and nL cells in the hindbrain. This dorso-lateral expression persisted and grew in size to HH st 30 when the beginnings of a demarcation of nM and nL cells could be detected. This expression of cadherin-2 in nM and nL persisted to at least HH st39. Additionally, the intracellular binding partner of cadherins, *γ-catenin* (also known as plakoglobin (Bello et al, 2012; Imamura et al, 1999) was also expressed within the nM and nL from at least HH st30 to HH st36 (data not shown). We next performed immunofluorescence using antibodies to cadherin-2 and γ-catenin to address the localisation of protein within the nM and nL. We found that both cadherin-2 and γ-catenin were not prominently localized to the soma of nM cells but instead predominantly localized to the bitufted dendrites of the nL (Figure 4P-O; Figure S6). Thus, cadherin family members are expressed in the nucleus laminaris and nucleus magnocellularis during the timecourse of nL formation as a lamina.

**Figure 4.**
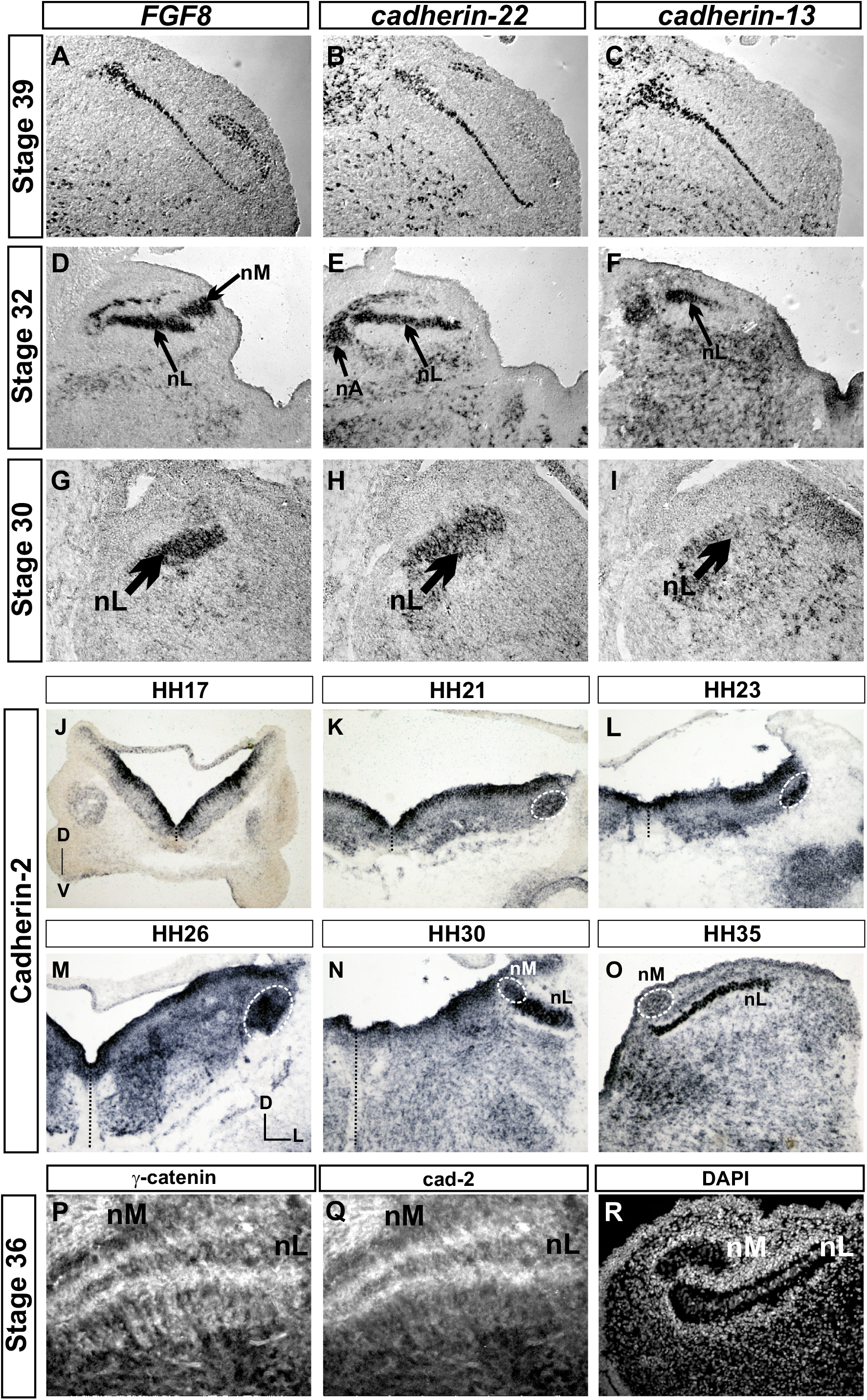
A-I *FGF8* (A,D,G), *cadherin-22* (B, E, H) and *cadherin-13* C, F, I) expression in the nM, nL and nA at HH stage 39, 32 and 30, as indicated in the left hand boxes. J-O *Cadherin-2* (*N-cadherin*) expression in the hindbrain at rhombomere 5/6 at HH stages 17 (J) 21 (K) 23 (L) 26 (M) 30 (N) and 35 (O) P-R. γ-catenin (P) and cadherin-2 (Q) expression in the nM and nL related to DAPI expression (R) at HH stage 36.

### An interplay between FGF signalling, cadherin expression and lamina formation in the auditory anlage

The timing of expression of *FGF8, MafB* and cadherin expression suggested that there may be a causal relationship between them. To address whether there may be an interplay between FGF8 expression, cadherin expression and lamina formation in the auditory anlage, we analysed expression of the four FGF receptor genes and Sprouty 4, a downstream target of FGF signalling (Dorey and Amaya, 2010; Labalette et al, 2011). Sprouty 4 has also been shown to be a negative-regulator of FGF signalling, indicating that it provides a negative feedback loop to tightly regulate FGF signalling. We found that FGF Receptors 1 and 2 and Sprouty 4 were expressed in the nL prior to and during lamina formation (Figure 5A-C), indicating that FGF signalling may be acting in an autocrine fashion within the nL.

**Figure 5.**
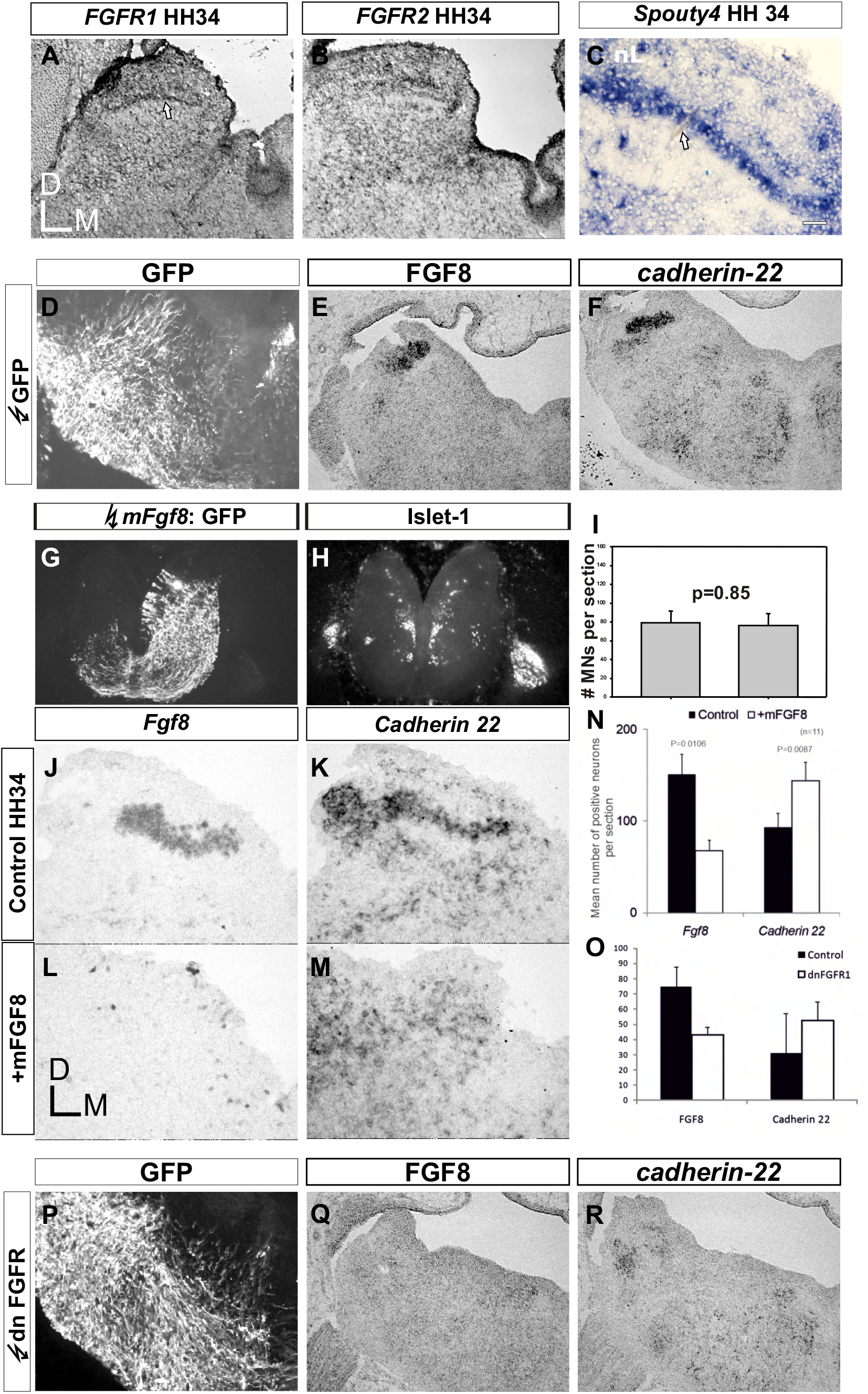
A-C *FGFR1* (A) *FGFR2* (B) and *Sprouty-4* (C) expression at HH stage 34 D-F control *in ovo* electroporation of GFP at HH st 18 followed by expression of GFP (D) *FGF8* (E) and *cadherin-22* (F) at HH stage 30. G-I mFGF8-GFP *in ovo* co-electroporation at HH st 18 followed by GFP (G) and islet-1 (H) expression at rhombomere 8 at HH stage 30. I Quantitation of motor neuron number at r8 of mFGF8 electroporation compared to control. J-M *FGF8* (J,L) and *cadherin-22* expression (K,M) following mFGF8-GFP co-electroporation or on the control side of the hindbrain as indicated in the left-hand boxes. N Quantitation of *FGF8* or *cadherin-22* cells in the nL area at HH st 34 following mFGF expression compared to control. O Quantitation of *FGF8* or *cadherin-22* cells in the nL area at HH st34 following dn FGFR1 expression compared to control. P-R dnFGFR-1-GFP co-electroporation at HH st18 followed by analysis of GFP (P), *FGF8* (Q) and *cadherin-22* (R) expression at HH stage 34.

We next sought to perturb FGF signalling through misexpression of mouse FGF8 (Fukuchi-Shimigori and Grove, 2001), or expression of a dominant negative isoform of FGF Receptor 1 (Saffell et al, 1997). Expression of a control plasmid expressing only GFP left unperturbed the formation of the nM and nL as a lamina, as assessed by FGF8 expression (Figure 5D-F). We next sought upregulation of FGF signalling through expression of mouse FGF8 via in ovo electroporation. We were concerned about potential developmental patterning effects, particularly in the AP axis, following FGF8 manipulation. Therefore, we sought a proxy for normal AP patterning in the organization and number of motor neurons in the caudal hindbrain. Following mFGF8 electroporation at HH st20 and analysis at HH st34, we detected no changes to either the number or the general organization of the glossopharyngeal (IXth), Vagal (Xth) or hypoglossal (XIIth) motor nuclei (Figure 5G-I). This gave us confidence to analyse formation of the nL following electroporation of mFGF8.

Following mFGF8 expression, we observed a decrease in the number of cells expressing chick FGF8 in the auditory anlage and also a perturbed nucleus laminaris structure formation as assayed by both cFGF8 and cadherin-22 expression (Figure 5J-O). FGF8 and cadherin-22 expressing cells of the auditory anlage appeared scattered and without the normal lamina-like structure.

We next asked what the effect of downregulation of FGF would be within the auditory hindbrain. We thus expressed a dominant negative DNFGFR1 expression construct and found a similar effect to upregulation of FGF8 expression. Following DNFGFR1 expression we observed a reduction in the number of cells expressing FGF8 and an abrogated auditory anlage formation as assayed by both FGF8 and cadherin-22 expression (Figure 5 O-R). Thus, there appears to be an interplay between FGF8 signalling, cadherin expression and lamina formation in the auditory anlage. Manipulation of FGF signalling both by upregulation and downregulation caused a disruption of lamina formation, as assayed by cadherin expression.

### Levels of FGF signalling are critical to nL formation

Our previous results suggest that levels of FGF8 signalling may be important to the formation of the nL lamina. To address this, we employed an in vitro slice-culture approach. During our initial attempts at culturing the brainstem and assessing lamina formation of the nL via FGF8 expression, we observed relatively few, scattered cells. We thus reasoned that any experiments utilizing this technique needed an automated approach to analysing nL lamina formation. To this end, we quantitated wild-type nL lamina formation via an automated algorithm. This algorithm identifies cells within the nL area and calculates for each section a coefficient of determination of the normalized average cell distance from a hypothetical lamina (see materials and methods). This r^2^ value approximates a numerical estimate of how linear the nL is-values closer to 1.0 being most linear, those closest to zero being the least linear (most voluminous). Utilizing this r^2^ calculation of wild-type FGF8 expression resulted in the values shown in figure 6A-F. Similar to what we observed before (Figure 3), the nL is least lamina-like at stage 30 (Figure 6A) and the major flattening of the nL into a lamina occurs from HH st33 to HH st36 (figure 6B-D; Figure 6M).

**Figure 6.**
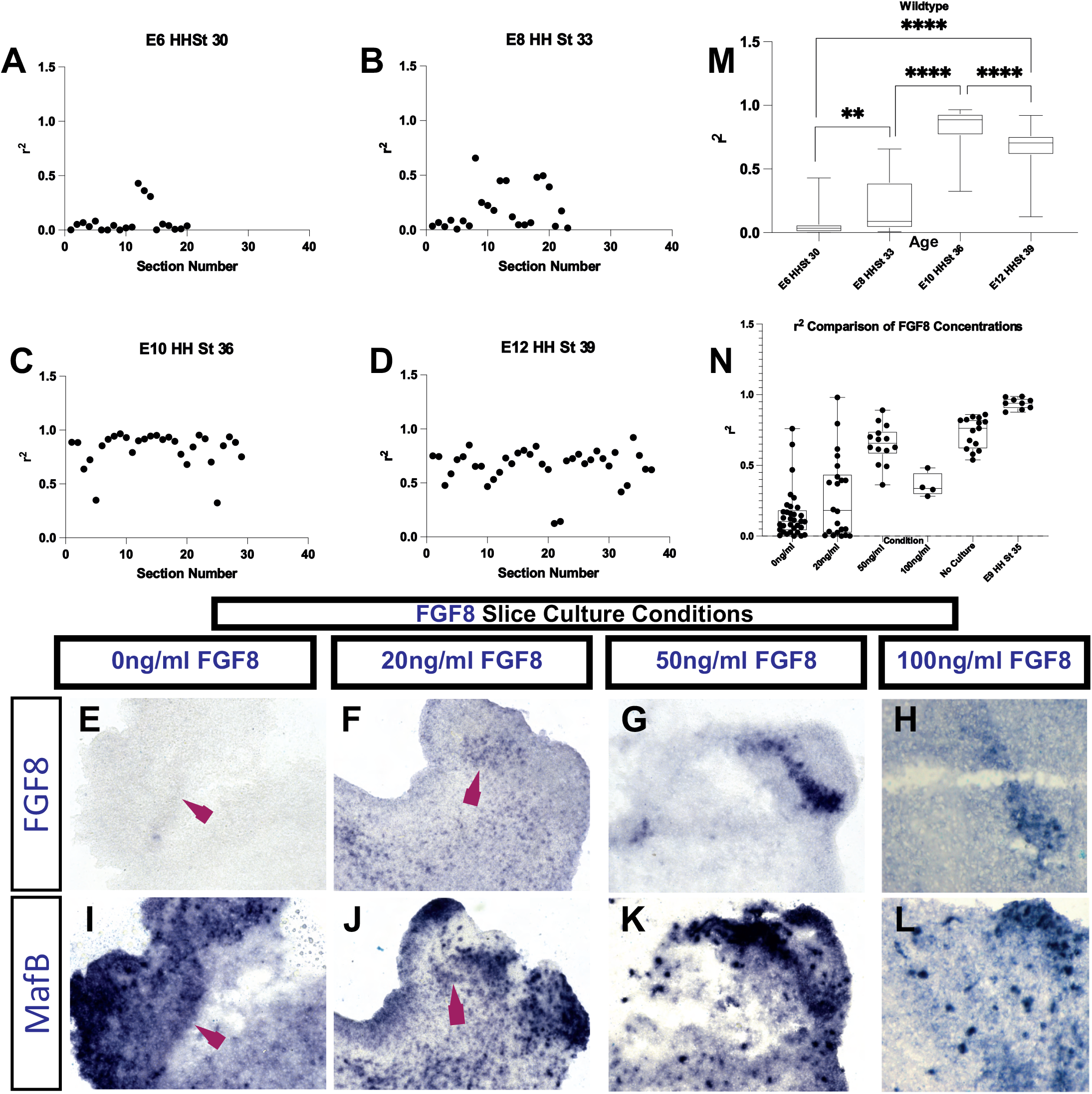
A-D r^2^ measures of lamina formation for each section of FGF8 in situ expression at HH stage 30 (A), st33 (B) st 36 (C) and st 39 (D) see materials and methods for details. E-L. *FGF8* (E-H) and *MafB* (I-L) expression in hindbrain slices cultured in varying concentrations of exogenously added FGF8 indicated in the boxes above). M, N Box and whisker plots of r^2^ calculations of wildtype at different HH stages and in different concentrations of FGF8 in culture conditions (N)

We thus sought to use the r^2^ measure of lamina formation to judge whether the addition of exogenous FGF8 to our slice cultures affected lamina formation as judged by FGF8 and MafB expression. Culturing brainstem slices for 24 hours from HH st34 in the absence of added exogenous FGF8 protein resulted in a severe downregulation of FGF8 expression in the slice (Figure 6E). Addition of 50ng/mL FGF8 protein during the culture period rescued the endogenous expression of FGF8 and MafB, along with lamina structure (Figure 6G, K). However, addition of either 20ng/mL or 100ng/mL FGF8 to the slice cultures mimicked the condition of no additional exogenous FGF8 protein (Figure 6F, J, H, L). Taken together, these results suggest that FGF8 expression in the nL signals within the nL and that levels of FGF8 signalling are important for lamina formation (Figure 6N).

### FGF8 regulates cadherin expression in the nL via MafB expression

We next asked whether the effects on lamina formation via manipulation of FGF signalling operate via MafB and cadherin expression. To test this, we assayed the expression of MafB following misexpression of FGF8. Following mFGF8 misexpression from HH st20 (Figure 7A-F), we found a dysregulation of MafB expression both at levels of the hindbrain that contain the auditory nuclei (dotted circles in Figure 7E, F) as well as in more rostral areas, where we observed increased expression of MafB (dotted circles in Figure 7B, C). We next asked whether MafB can regulate cadherin expression within the brainstem. We first misexpressed MafB and assayed MafB expression itself in the auditory hindbrain. Following MafB expression, we confirmed an increase in expression of MafB cells in areas directly adjacent to the auditory hindbrain (Figure 7 G-I). In these areas displaying ectopic MafB expression, we also detected ectopic cadherin-22 expression (Figure 7J-L). This suggests that MafB expression can drive cadherin-22 expression. We next attempted to downregulate MafB function through the expression of a dominant negative MafB construct (Figure 7M-R). Following this expression (figure 7M-R), in areas where dnMafB expression was detected, we also detected a downregulation of cadherin-22 expression within the auditory hindbrain (dotted area in Figure 7Q). In both of these conditions, we were unable to detect a change in expression of cadherin-13 within the auditory hindbrain, indicating that MafB probably does not control the expression of cadherin-13 (data not shown).

**Figure 7.**
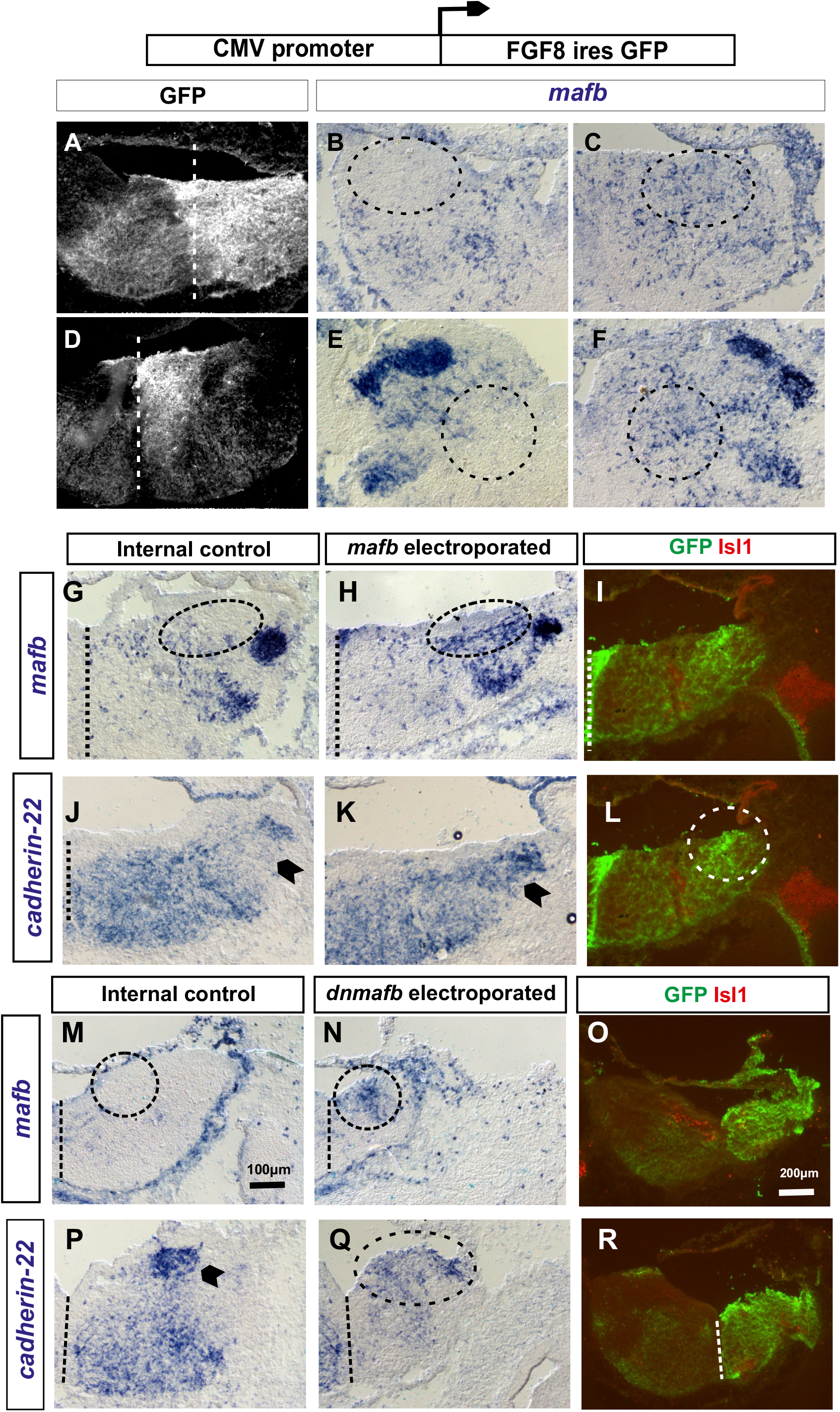
A-F. GFP (A,D) and *MafB* (B,C,E, F) expression following in ovo electroporation of mFGF8. Analysis was at HH stage 30 in r3 (B, C) and r6 (E, F) for control side of the hindbrain (B, E) and experimental side (C, F). G-L *mafB* (G-I) and *cadherin-22* (J-L) expression following in ovo electroporation of MafB. Internal control side of the hindbrain (G, J) and experimental sides (H,K, I, L) are shown. electroporation is marked by GFP (I, L) Isl-1 expression shows location of r5 motor nuclei and ganglia. M-R. dnMafB electroporation is detected by either GFP (O, R) or MafB experssion (M,N). P, Q shows its effect on Cadherin-22 expression.

However, these results together suggest that the actions of FGF8 expression within the auditory hindbrain could, at least in part, be mediated via MafB expression within the auditory hindbrain controlling cadherin-22 expression.

### Cadherin function is critical to the lamina formation of the nL

We next asked whether cadherin function was critical to the formation of the nL as a lamina structure. We first asked whether overexpression of cadherin-2 impacts lamina formation. Following expression of cadherin-2 via a doxycycline inducible plasmid (Tanabe et al 2006) (electroporation at HH st19 and dox induction at HH st30), we detected no change in nL formation as a lamina as assessed by FGF8 expression at HH st36 (Figure 8A, B). We next assessed the effect of downregulation of cadherin function by expression of a dominant negative isoform of cadherin-2 within the auditory hindbrain. This dominant negative has been shown to dysregulate cadherin function through uncoupling cadherins from the actin cytoskeleton (Fujimori and Takeichi, 1993). As lamina formation occurs relatively late in embryogenesis, we sought to control the prolonged expression of the dominant negative isoform through the use of the doxycycline inducible transposase integrated version (Tanabe et al., 2006). Expression of NΔ390 via electroporation at HH st19 with induction of expression at HH st30 was performed with visualisation of FGF8, cadherin-22 and cadherin-13 expression at HH st36 in rostral nL which, by this stage is flattening into a lamina structure. Following NΔ390 induction, cadherin-22 and cadherin-13 expression on the electroporated side showed that nL formation as a lamina was perturbed (Figure 8C-F); compared to the control side of the embryo, NΔ390 expression left nL formed as a ball of cells, similar to that found in the early auditory anlage (Figure 8 D, F). This suggests that cadherin function is critical to the formation of the lamina of the nL. However, we also observed a striking downregulation of FGF8 expression in the nL and nM cells following NΔ390 expression (Figure 8E). Taken together, this suggests that downregulation of cadherin function causes a disruption of normal lamina formation both through a disruption of the organization of the cells as well as in FGF8 expression.

**Figure 8.**
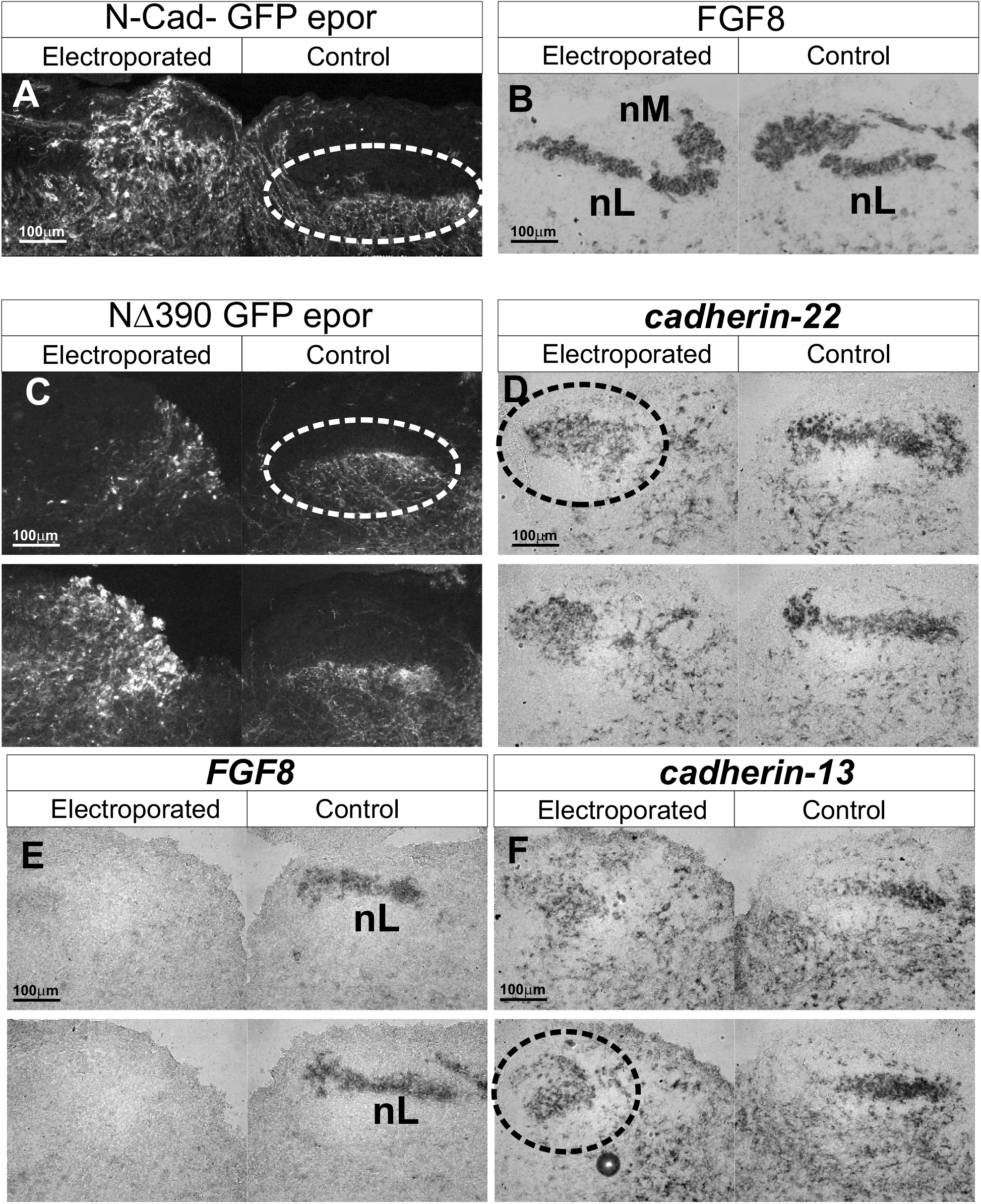
A-F In ovo electroporation at stage 19 of either N-cadherin (A, B) or dominant negative N-cadherin (C-F) doxicycline was added at HH stage 30 and analysis was at HH stage 36. GFP (A, C) marks electroporated cells and axons. Analysis of nL/nM is shown by *FGF8* expression (B, E), *cadherin-22* (D) or *cadherin-13* expression (F). Experimental or control sides of the hindbrain are shown as indicated in the boxes above.

### Modelling suggests that cell patterned cadherin function could drive lamina formation in the nL

Taken together, our results suggest that FGF and MafB regulated cadherin expression plays a critical role in the lamina formation of the nucleus laminaris. To address how cadherin expression could drive the flattening of a cell assembly into a flat sheet, we modelled the adhesion effect of cadherins on the cell population configuration, taking into account that cadherin expression was observed to be localized on bipolar nL dendrites (Figure 4Q, Fig S6).

As a simplified, ‘toy’ model, each cell was considered as a square-based rod unit in a close-packed lattice arrangement (Fig 9A) and the overall structure of the cell population was assumed to tend towards a maximum adhesion energy configuration. In the case of purely adhesive forces between cells, this is also the maximum adhesion configuration. In this simple model, adhesion between any two units is proportional to their contact area and the total contact area is dependent on both the population configuration and the unit structure (Fig 9B). To solve the configuration of the assembly that maximises the total adhesion energy of the system, we imposed a constraint on the model that the total number of cells remain constant. We reasoned that this is a good approximation to the early beginnings of formation of the nL as a lamina in rostral regions of the nL prior to the peak of apoptosis. It is also assumed that interactions between the outermost cells of the assembly and surrounding non-nL cells have lower adhesion than those between nL cells, as otherwise the nL cell group could dissociate.

**Figure 9.**
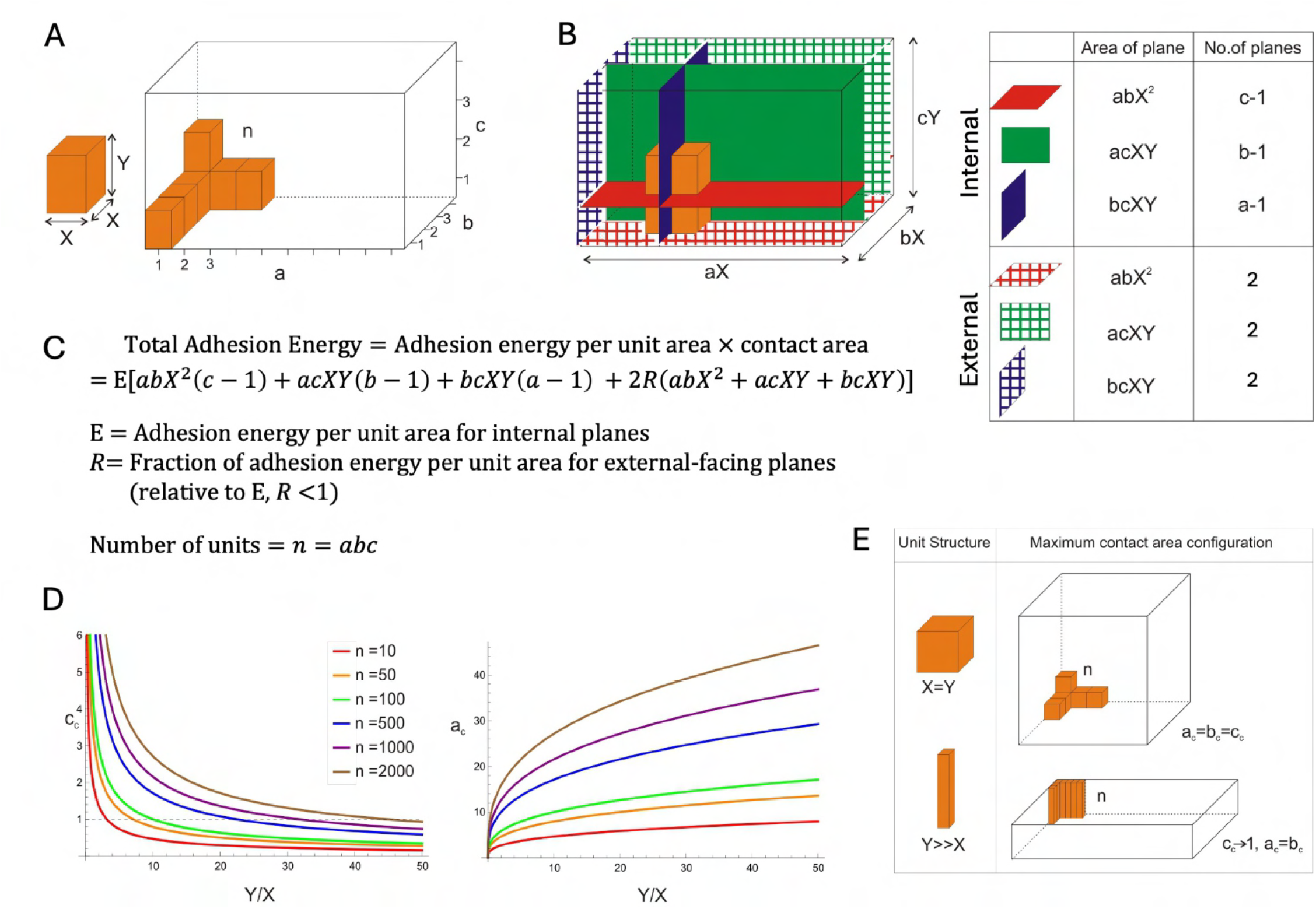
(A) Diagram of simple square-based rod model for n units. Square-based rods have width X and height Y and are arranged on an orthorhombic lattice. There are a total of n units, with a, b and c integer units along the x, y and z axes respectively. (B) There are three different types of contact planes between adjacent units: x-y planes (red), x-z planes (green) and y-z planes (blue). The number and area of each type of plane are shown for internal planes (between units in the population of interest) and external planes (between the population of interest and surrounding media). (C) Outline of total adhesion energy calculation for a constant number of units (n) with uniform adhesion per unit area for internal contact planes and a reduced adhesion per unit area for external facing planes. (D) Dependency of the population maximum adhesion configuration (a_c,_ b_c,_ c_c_) on the individual unit structure. Solutions for the constrained maximum (Appendix 1) have critical values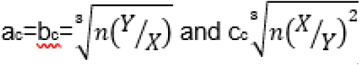 The minimum possible values of a_c_ and c_c_ are 1 as a, b and c take integer values. (E) The maximum adhesion configuration for a cubic unit structure (Y/X = 1) is cubic. Tall, narrow, unit structures (Y/X >> 1) show a flattened maximum adhesion population.

Using the method of Lagrangian multipliers to solve for constrained critical values (Appendix 1), we find that the maximum adhesion configuration depends on the width:height ratio (X:Y) of the cell unit and the total number of cells (n). When the X:Y ratio is 1:1 (cubic unit structure) the population has a maximum adhesion configuration which is also cubic but as the unit structure becomes taller and narrower the maximum adhesion structure becomes increasingly laminar (Fig 9C).

This ‘toy’ model highlights how the spatial adhesive characteristics of individual cells impacts the spatial configuration of the cell group. To extend the generality of these findings, a more abstract model was developed that considers the cells as interacting points on a finite lattice structure (Appendix 2). Instead of considering contact areas, interactions between adjacent points are considered as adhesion energies along primitive lattice vector directions. In this more general model, the maximum adhesion configuration of the cell group is dependent on the ratio of adhesion energies along the primitive lattice vector directions (shown for orthorhombic and triclinic lattice structures). Specifically, when adhesion energy along one axis is lower than along the others, the maximum adhesion configuration has a reduced extent in the direction of lower energy.

These models are consistent with adhesion in the dendritic domain of each nL cell being able to drive lamina formation of the nL owing to this configuration maximising the total adhesion energy of the system. Thus, in both a more generalized model of cell-adhesion displaying a preferential axis and in a simpler square-based rod model of adhesion, both models predict that lamina formation results in the maximum adhesion configuration for a group of cells with bipolar morphology.

Our previous models considered the static, maximum adhesion configuration of cells. We next sought to model lamina formation via a more dynamic model (Fig S7). We considered and modelled each nL cell as comprising bitufted adhesive units built on springs capable of moving in two dimensions (Fig S7A, B). We then modelled different numbers of such cells within a defined box of fixed dimensions (Fig S7C).

Cells were initially localized at random places within the box and then the simulation allowed cells to move freely within the box. Following rounds of dynamic simulation, various adhesive factors were refined (Fig S7D, E). We observed that in these dynamic simulations, equal adhesion between both “dorsal” versus “ventral” adhesive units resulted in clusters of cells, more ovoid in shape (Fig S7E). However, if we modelled the dorsal adhesive unit to preferentially interact with dorsal units versus ventral units preferentially interacting with ventral units then the maximum adhesion configuration resulted in a more lamina-like structure forming (Fig S7D). Taken together, both static as well as dynamic models of the nucleus laminaris that take into consideration a cell-patterned, bipolar morphology of adhesive units suggest that lamina formation is the adhesively most favorable configuration.

## Discussion

In this study, we demonstrate expression of FGF8 and members of the FGF signalling pathway along with members of the cadherin family within the developing auditory hindbrain of the chick. We show expression within the nucleus laminaris during lamina formation. Disruption of FGF signalling perturbs cadherin expression and lamina formation. We further show that FGF expression can regulate MafB expression and that MafB expression can regulate cadherin-22 expression. Cadherin expression is localized within the dendrites of the nL and we both show and model how cadherin function could drive formation of the nL cells as a lamina. We discuss these findings in the context of a model whereby levels of FGF signalling within the nL regulate cadherin expression, partly via MafB expression, which drives lamina formation of the nucleus laminaris.

Several lines of evidence suggest that FGF8 functions in an autocrine fashion within the nL. First, FGF8 expression is prominent in the auditory hindbrain during later embryogenesis. There is very little expression of FGF8 within the hindbrain from stage 30 to stage 39 other than that in the nM and nL. Additionally, FGF receptors are expressed in the nL during lamina formation and sprouty-4, known to be a downstream target of FGF signalling is also expressed within the nL. Second, misexpression of mouse FGF8 within the chick hindbrain perturbs the endogenous expression of FGF8 within the auditory hindbrain. Third, in vitro hindbrain slice cultures require supplements of purified FGF8 protein in order to facilitate lamina formation and FGF8 expression. We show that levels of added FGF8 are important, with 50ng/mL being optimal in our culture conditions. Sprouty-4 is known to be a negative regulator of FGF signalling and thus could provide a source for a negative feedback loop to ensure that levels of FGF signalling are maintained or tightly controlled. Autocrine loop signalling with FGFs have been described in the literature (Park et al, 2008), albeit outside the nervous system, providing precedence for such an effect in the hindbrain.

We also show that FGF8 signalling controls the expression of MafB expression within the hindbrain, suggesting that MafB may also be a downstream effector of FGF signalling within the auditory hindbrain. FGF signalling has also been shown to regulate MafB expression in other regions of the nervous system. For example, FGFs regulate both Krox-20 and MafB expression in the otic/pre-otic region of the chick hindbrain (Marin and Charnay, 2000). Additionally, sprouty-4 has been shown to regulate krox-20 signalling in the hindbrain (Labalette et al, 2011), thus further making a potential link between levels of FGF signalling and the MafB/ Krox-20 family member expression.

We also show that MafB expression can regulate cadherin expression, suggesting that a role for FGF8 regulated MafB expression might be the expression of cadherin-22 within the nL. Relatively little is known about the cadherin-family members that may be downstream of the MafB transcription factor. However, MafB has been shown to regulate E-cadherin (cadherin-1) in lens formation, indicating that there is precedent for MafB regulation of cadherin genes, if not cadherin-22 (Reza et al 2007).

We further demonstrate that cadherin-2 (N-cadherin), cadherin-13 and cadherin-22 are expressed within the nucleus laminaris, with cadherin-2 potentially demarcating cells of the auditory hindbrain shortly after they are born. Combinatorial cadherin expression, including that of cadherin-2, a type I cadherin, with members of the type II family of cadherins have been shown to regulate cell-patterned clusters of soma of neurons in multiple areas of the nervous system, notably within spinal cord and brainstem motor neurons (Astick et al., 2014; Price et al., 2002; Egusa et al., 2016; Vagnozzi et al., 2022). For example, differential cadherin expression in spinal motor neurons controls the clustering of motor pools that project axons to different peripheral muscles. Cadherin function also controls the migration of those motor neurons to the ventral horn (Bello et al., 2012). Additionally, combinatorial cadherin expression controls the clustering of brainstem motor somatic nuclei including those that project axons to the extraocular eye muscles. A similar combinatorial cadherin expression profile has also been observed to control the clustering of phrenic motor neurons that control breathing (Vagnozzi et al., 2022). In that location, this cadherin profile also controls the dendritic architecture of the phrenic motor neurons. Dendritic architecture has also been shown to be controlled by cadherin expression in the mouse primary somatosensory barrel field (Egusa et al., 2016). This connection, we posit is significant in so far as it demonstrates a link between cadherin expression and function and neuronal dendrite architecture. This suggests that dendritic cadherin function may also play a causative role in the patterning of dendrite architecture. Our results suggest that cadherin function is important to the formation of the nL as a lamina of cells. First, expression of a dominant negative cadherin construct perturbs the normal lamina formation. Second, we show that cadherin along with intracellular gamma catenin expression is predominantly localized to the bipolar dendrites of the nL. Third, we show that a bipolar configuration of adhesive contacts is sufficient to drive an assembly of cells towards a lamina structure.

Importantly, our dynamic model of nL formation required us to incorporate a differential adhesion between dorsal versus ventral adhesive domains. Members of the ephrin and eph family of receptor tyrosine kinases, which have been shown to also elicit an adhesive effect, have been shown to be differentially expressed between dorsal and ventral dendrite domains within the nL with perturbation studies showing effects on tonotopic projections in the auditory brainstem (Person et al., 2004; Huffmann and Cramer, 2007). Whilst we have not been able to show such a difference in expression of cadherins, largely owing to the paucity of antibodies to cadherin family members that work in thin tissue section immunofluorescence, our results, together with the expression of Ephrin/eph, suggest that differential adhesive properties may underpin lamina formation of the nL.

Is there evidence for dendrite-based adhesion promoting cell assembly pattern and formation? Cadherin function has been shown to regulate the lamina synapse specificity of retinal axon targeting and, within the eye, dendrite fields of retina bipolar cells have a “tiled” morphology that indicates that dendrite interactions may underpin that patterning (Basu et al., 2015). Further, within the spinal cord, motor neuron pools that are located more centrally within the ventral horn tend to have radially symmetric dendrite fields and more ovoid soma clustering modes. However, those motor neuron pools that are located more peripherally within the ventral horn tend to display more tangential or curved dendritic fields with more crescent shaped clusters of soma (Vrieseling and Arber, 2006). The shape of these dendritic fields is thus related to the shape of the cluster of motor neuron soma. These variable dendritic field morphologies have been shown to be critical to the positioning required for appropriate synaptic input to motor neurons from 1a proprioceptive sensory neurons, thus indicating that they may be critical to motor neuron physiology (Balaskas et al., 2019). We thus suggest that there is precedent for cell-patterned, dendrite-based, adhesion activity by cadherin family members could drive the shape of the nucleus laminaris as a lamina.

What may be the consequences of dendrite-patterned adhesion in lamina formation in the nL? During auditory hindbrain development, cells within the auditory anlage, particularly the nL, have been shown to elaborate bipolar dendrites in random directions, initially uniformly throughout the nL. These bipolar dendrites subsequently refine in architecture such that the nL cells found most rostromedially, which will process higher frequency sounds, elaborate short, highly branched dendrites. In contrast, nL cells more caudolaterally, which will process more low-frequency sounds, elaborate longer, less branched dendrites. This dendrite architecture is thought to underpin the ability of nL cells to act as coincidence detectors for synaptic input from both ears and thus critical to sound source localization (Agmon-Snir et al., 1998). The juxtaposition of nL cells within the lamina is thus critical to the frequency domain of sound source localization in addition to being critical to the detection of interaural time differences by coincidence detection. Therefore, a cell-patterned, dendrite-based adhesion mechanism that was critical to the formation of the nL as a lamina could underpin these critical features of segregated frequency coding as well as appropriate representation of auditory space within the nucleus laminaris.

## Acknowledgements

We thank Zahra Samji for her initial work on the project. M.C. was supported by a BBSRC PhD student, M.A. was supported by an MRC PhD studentship and K.T. by a Wellcome Trust PhD studentship. This research was funded in whole, or in part by the Wellcome Trust [Grant number 094399/B/10/Z]. Data will be made available in accordance with funder guidelines.

---

**Fig S1.**
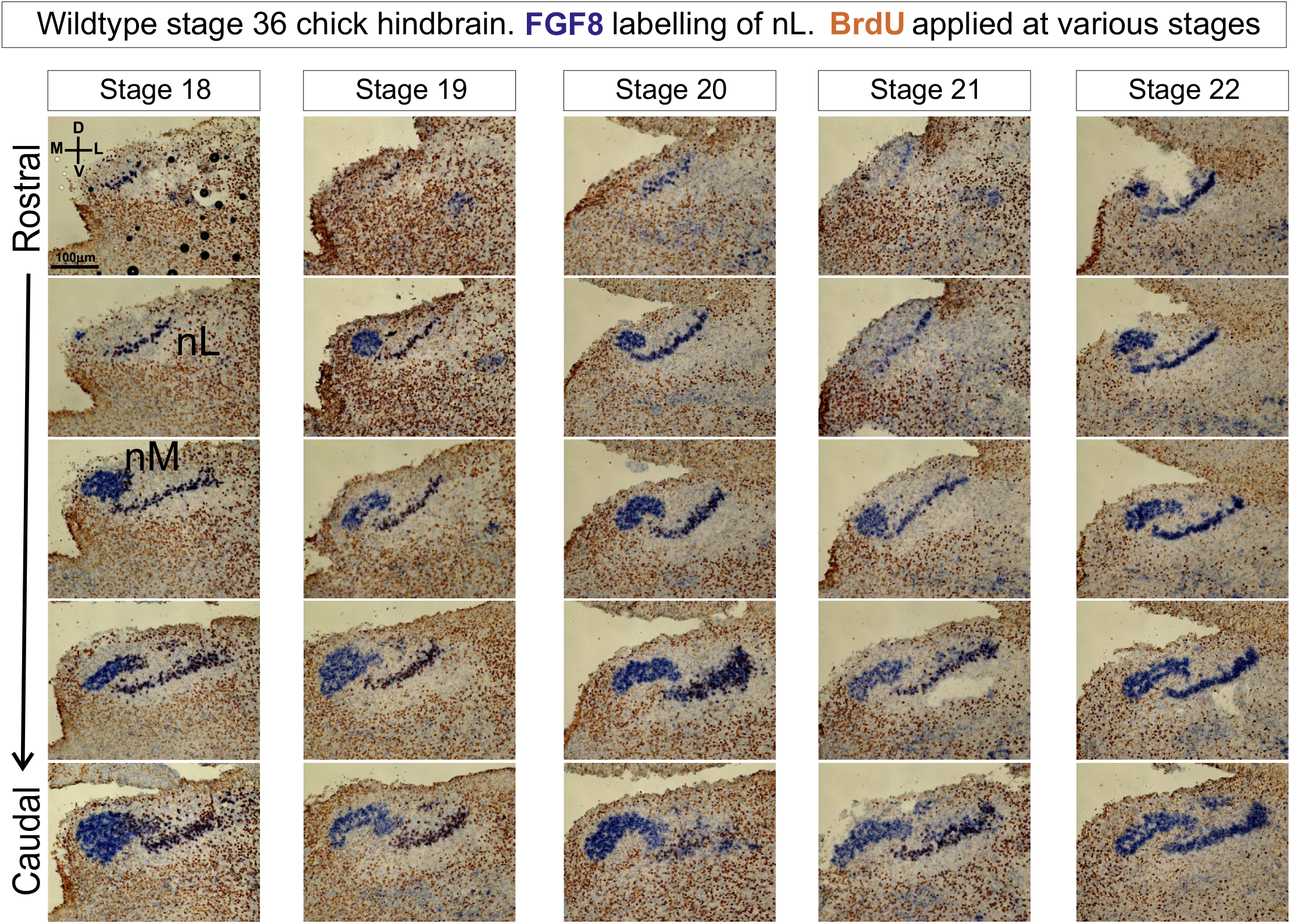
FGF8 in situ hybridisation at HH stage 36 following BrdU application to embryos at HH stages 18 to 22, as indicated by the boxes above the panels A series of sections from individual embryos is shown from rostral to caudal (top panels to bottom panels) as indicated by the schema on the left hand side.

**Fig S2.**
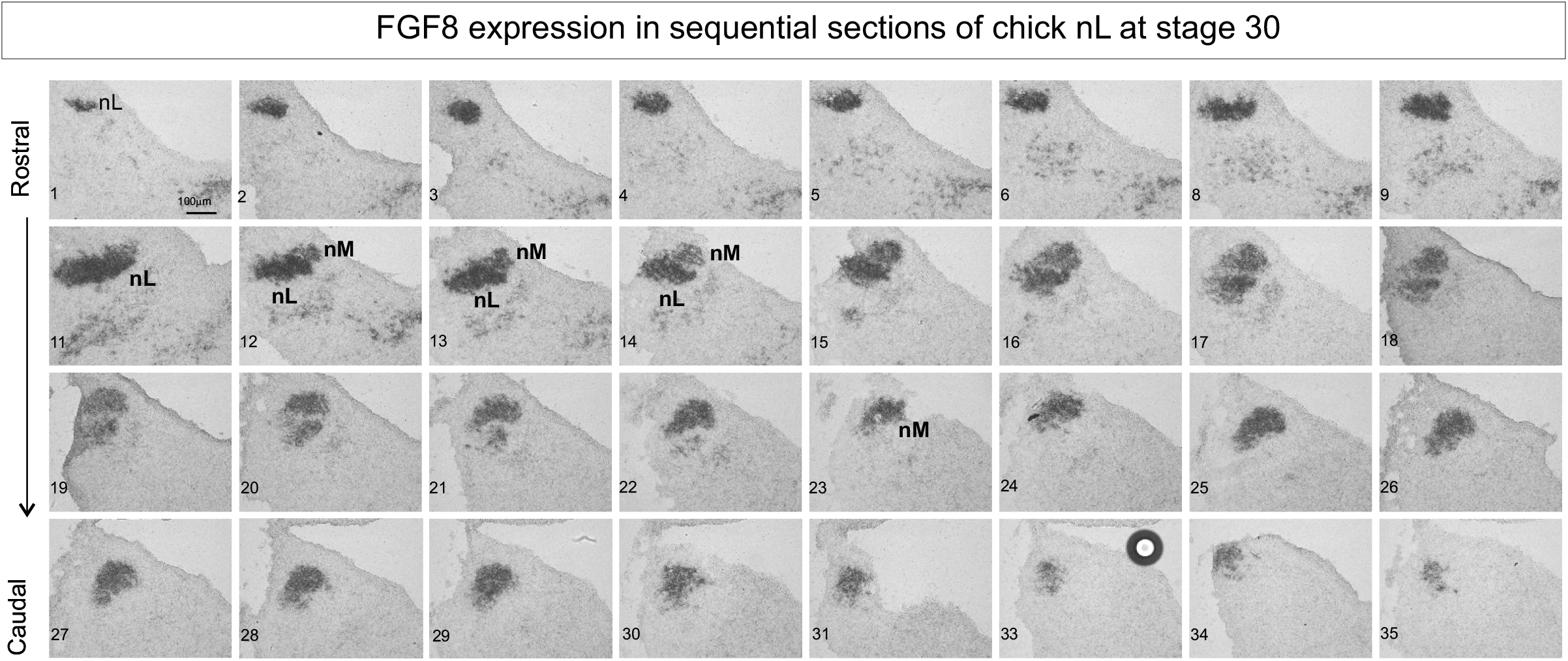
FGF8 expression at HH stage 30 across the entire auditory hindbrain sections are numbered sequentially from rostral to caudal most and are indicated in the panels.

**Fig S3.**
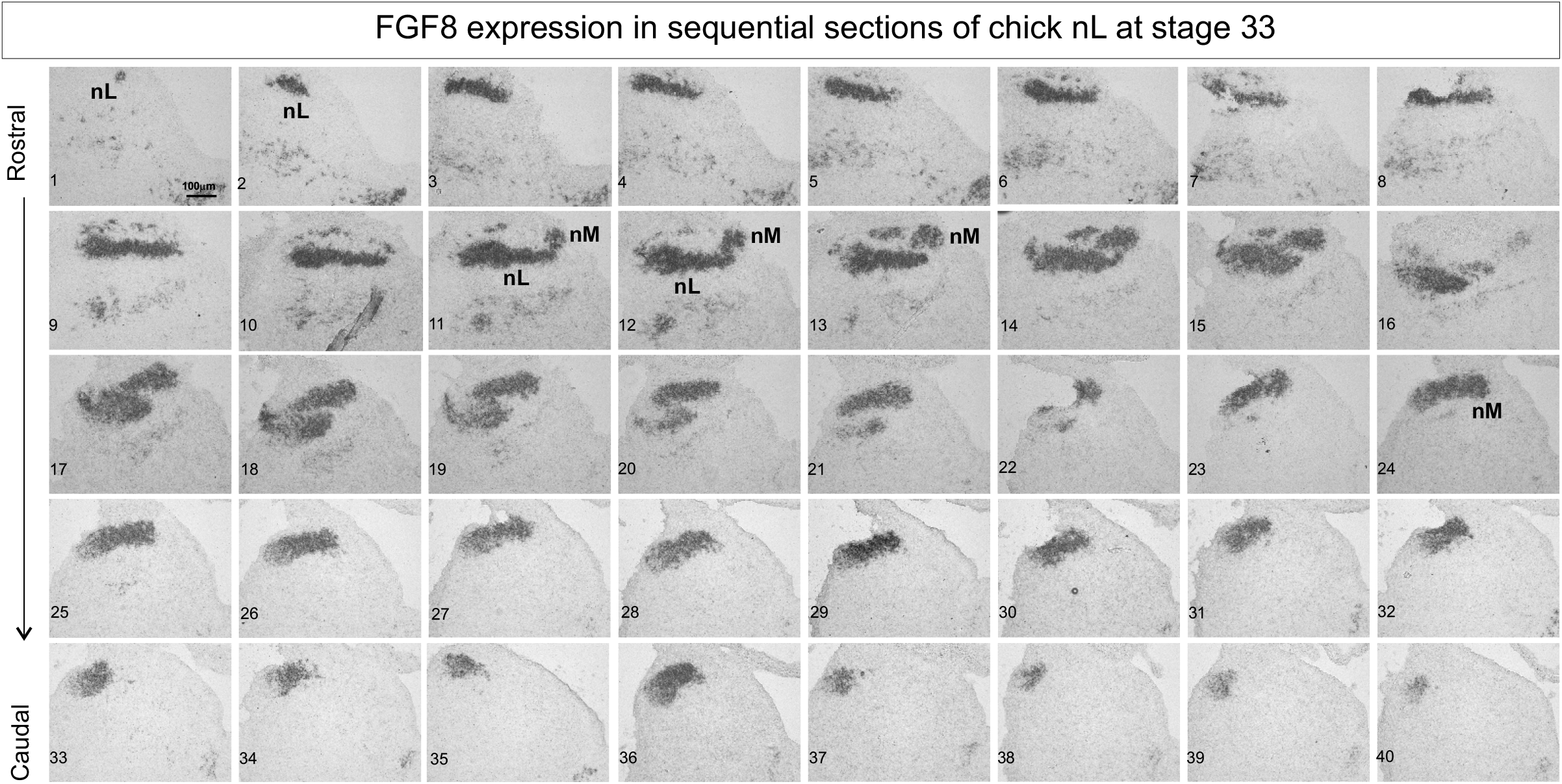
*FGF8* expression at HH stage 33 across the entire auditory hindbrain sections are numbered sequentially from rostral to caudal most and are indicated in the panels.

**Fig S4.**
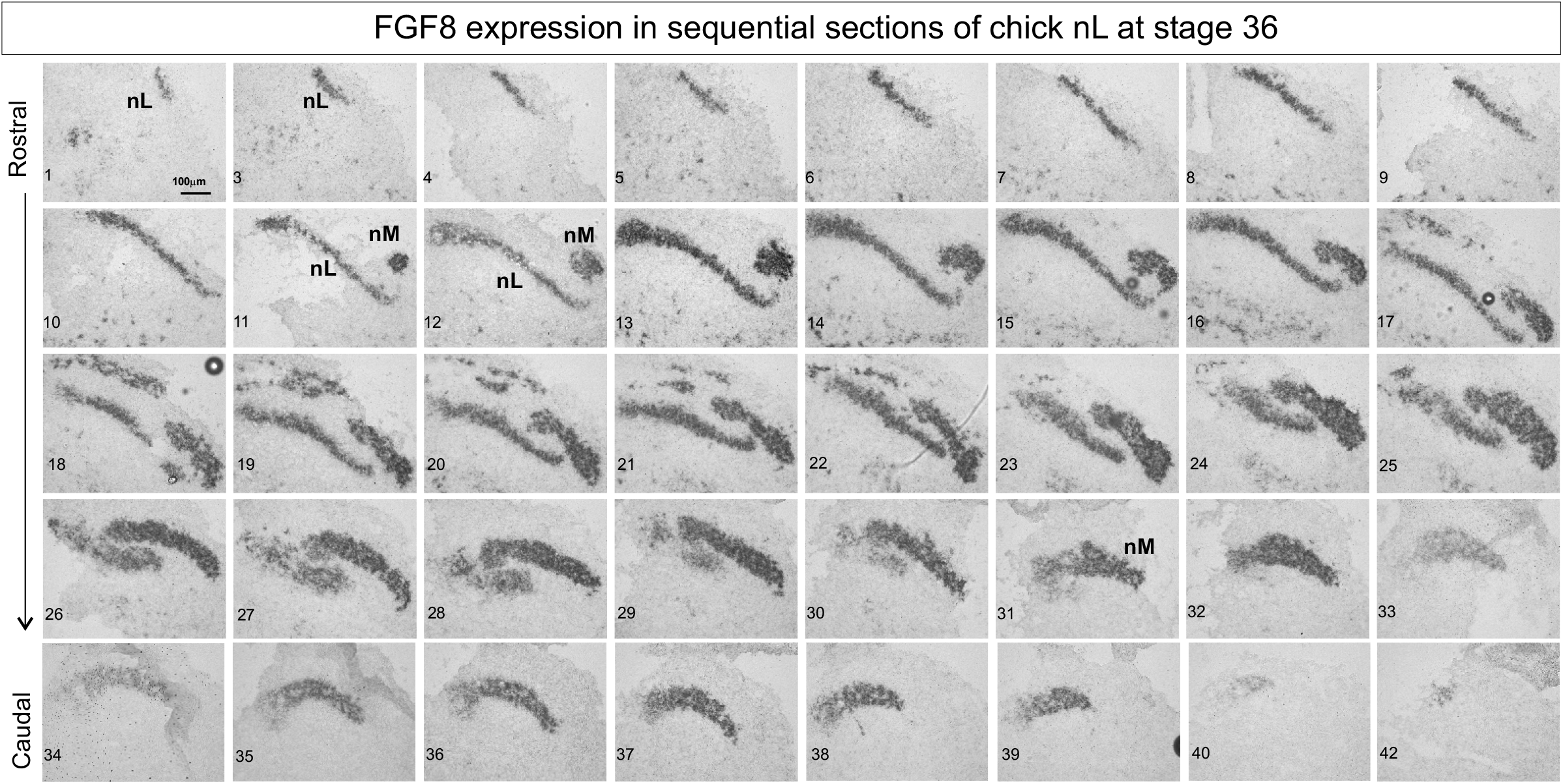
FGF8 expression at HH stage 36 across the entire auditory hindbrain sections are numbered sequentially from rostral to caudal most and are indicated in the panels.

**Fig S5.**
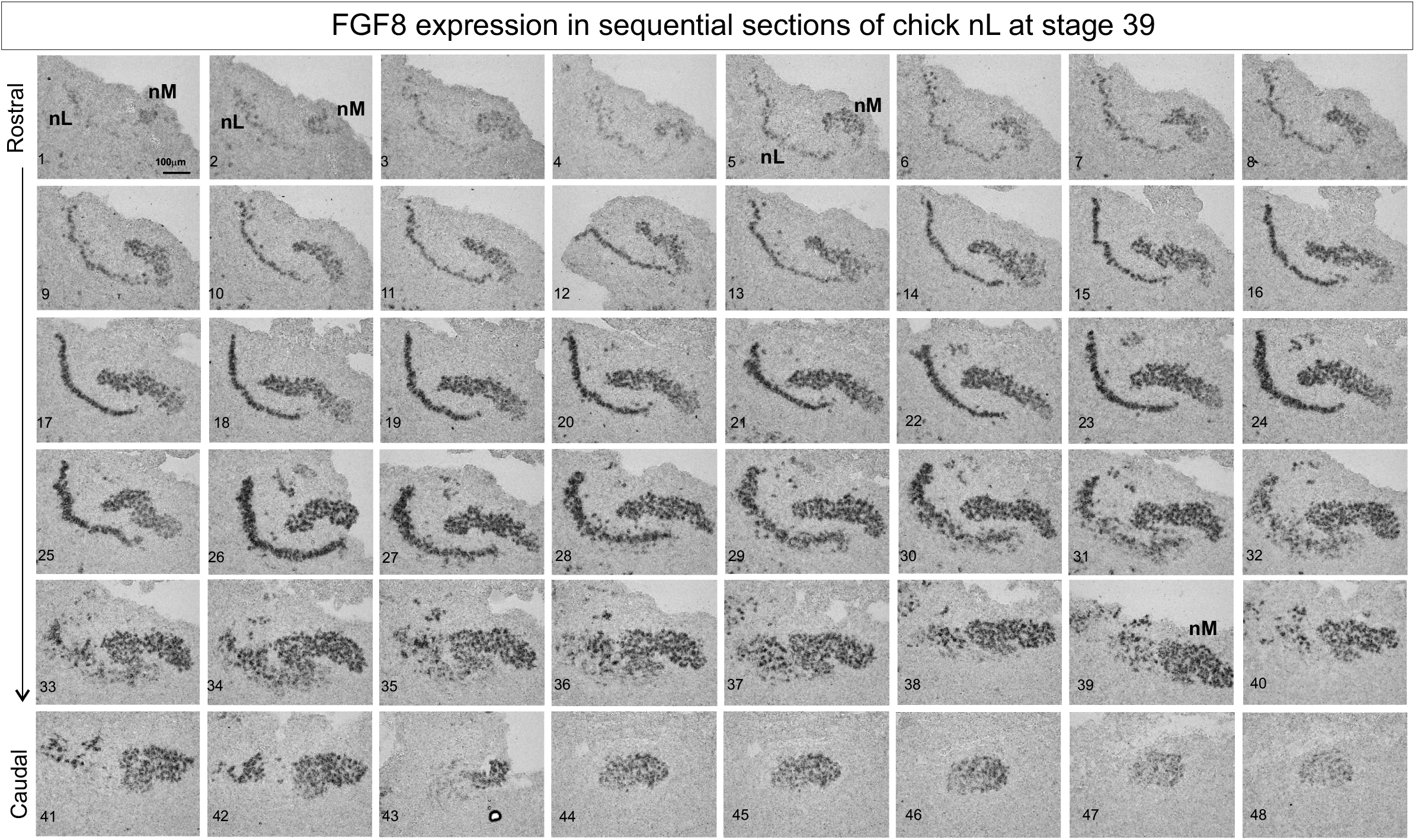
FGF8 expression at HH stage 39 across the entire auditory hindbrain sections are numbered sequentially from rostral to caudal most and are indicated in the panels.

**Fig S6.**
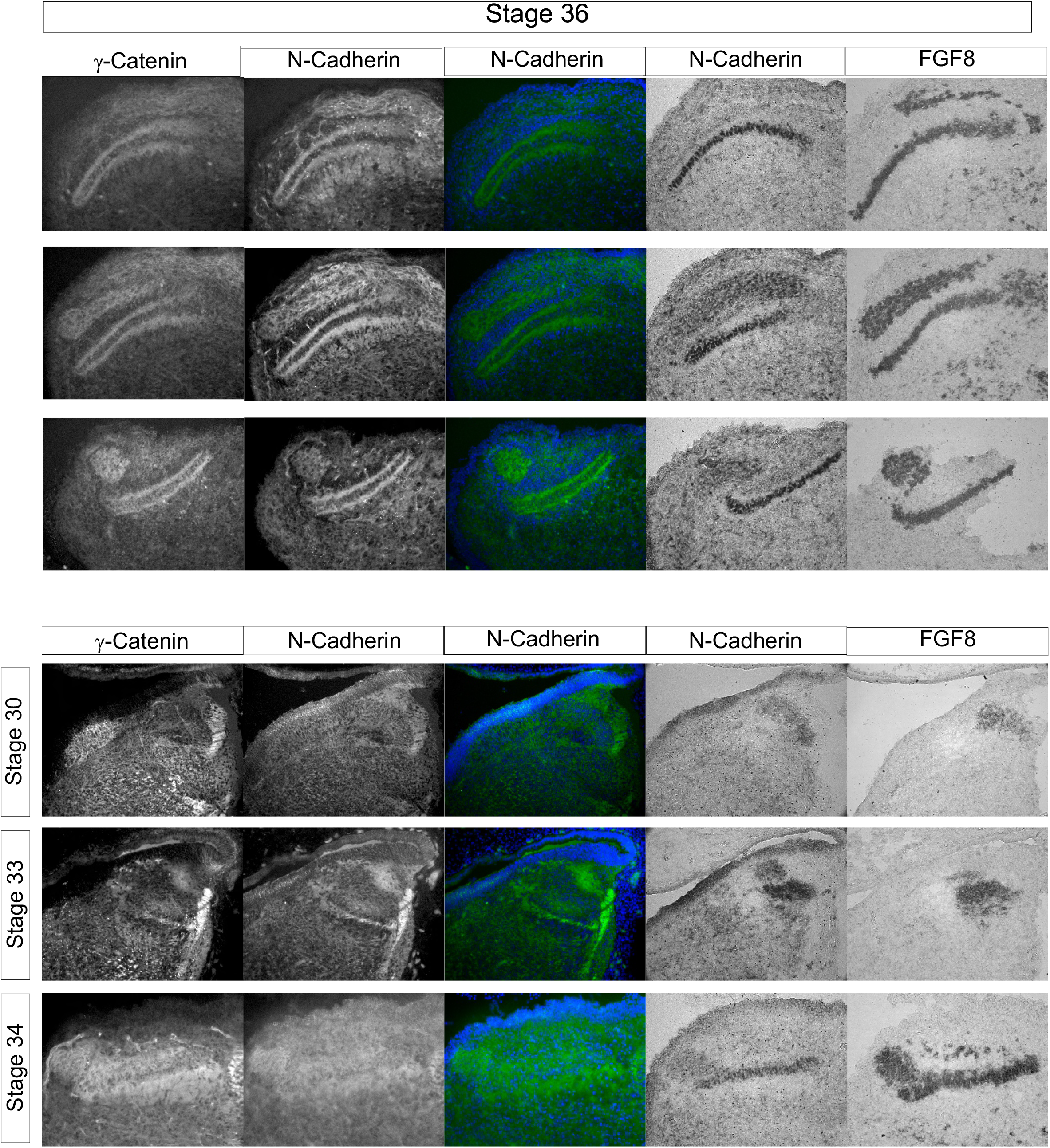
γ-catenin, N-cadherin, *N-cadherin* and *FGF8* expression at HH stage 36 top panels Various sections across the auditory hindbrain are shown. Bottom panels show a time series of expression in the auditory hindbrain at HH stage 30, stage 33 and stage 34.

**Figure S7.**
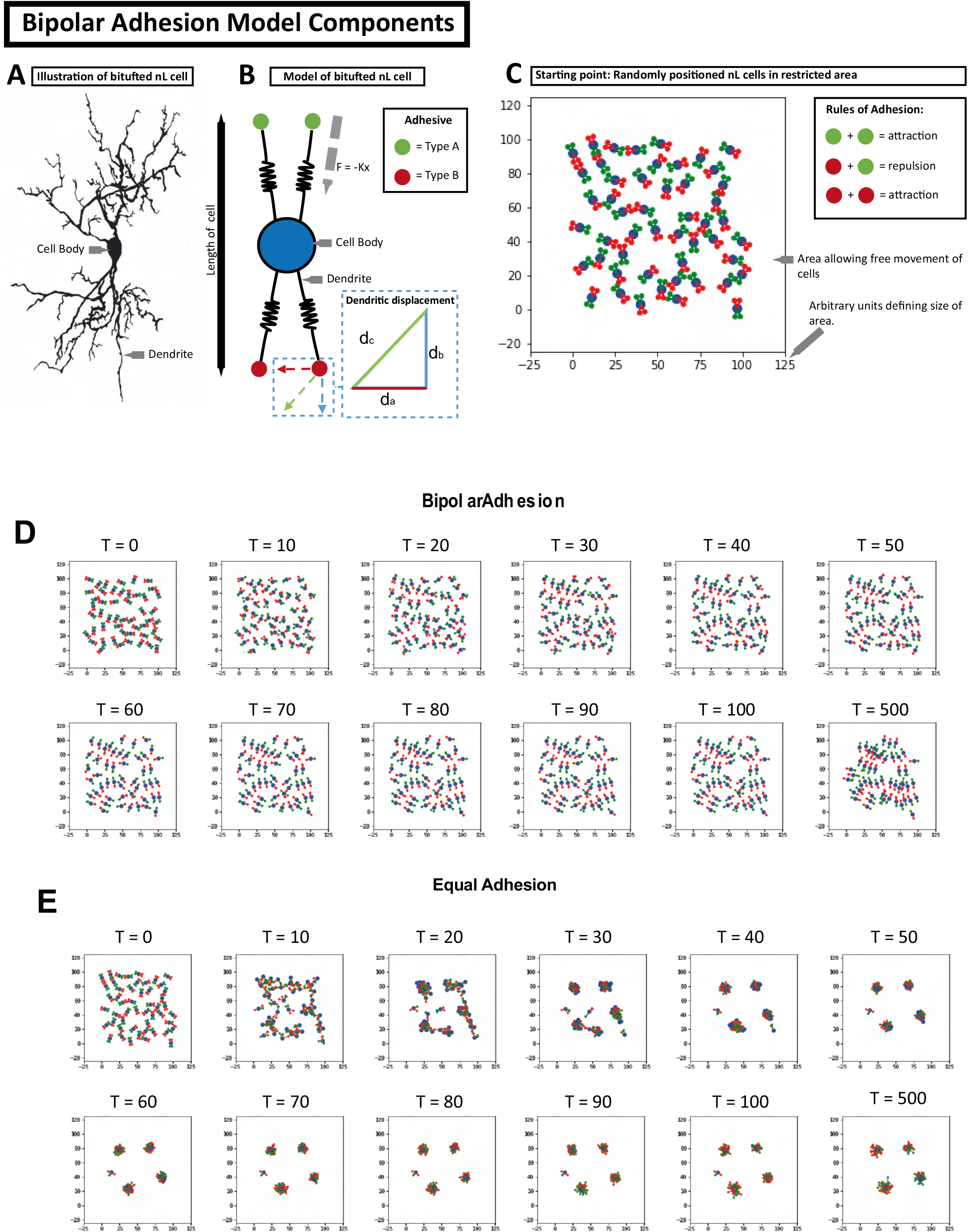
A-C. Dynamic adhesion model for nL formation.A. Schematic of a typical nL cell from the literature. nL cells show a characteristic bipolar morphology with dorsal and ventral dendrites. B. Schematic of the modelling of nL cells with a cell body and a dendrite modelled as springs capable of synamic displacement. C. Starting conditions of the dynamic model (see materials and methods) D. Time series ofthe dynamic model run with conditions of differential dorsal and ventral adhesion (bipolar adhesion conditions). E. Time series of the dynamic model run with conditions of equal adhesion potential between dorsal and ventral dendrite compartments.

## Appendix 1

This appendix expands on the square-based rod model of the nucleus Laminaris (nL) described in the main paper. In summary, nL neurons with adhesive bipolar dendritic fields are modelled as square-based rod units in a configuration of *a* units wide, *b* units deep and *c* units high, where the square-based rod is *X* wide and *Y* tall (Figure 9). The model considers the configuration that maximises the total adhesion energy of the square-based rod population. Adhesion energy within the square-based rod population is calculated as the total internal contact area between units multiplied by an adhesion energy per unit area (*E*). Adhesion energy with external surrounding media is calculated as the total external surface area multiplied by *ER*, where *R <* 1 is the ratio of internal to external adhesion. The total adhesion energy is given by:

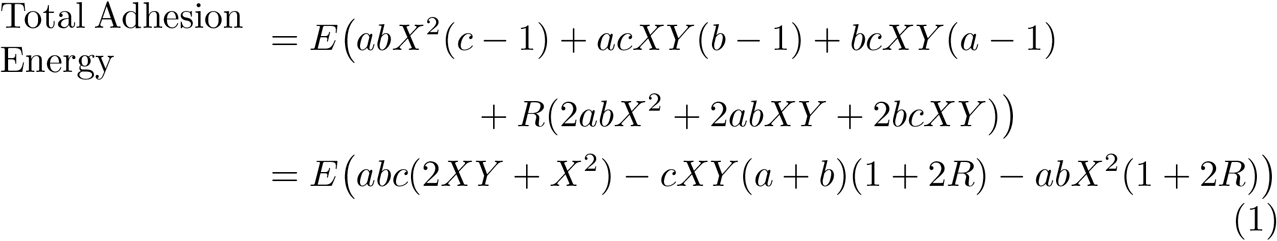

We wish to find the configuration (the values of *a, b* and *c*) which provides the maximum total adhesion energy for given values of X and Y given the constraint of a finite number of units (*n*). To find the critical points (maxima, minima or saddle points) of a function subject to a constraint, the method of Lagrangian multipliers [1, 2] is used. For a three-dimensional function *f* (*a, b, c*) subject to the constraint *k*(*a, b, c*), critical points exist where the vector gradients of *f* (*a, b, c*) and *k*(*a, b, c*) are parallel (equation 2). *λ* is an arbitrary constant, indicating that the condition requires the gradients to be parallel, not equal.

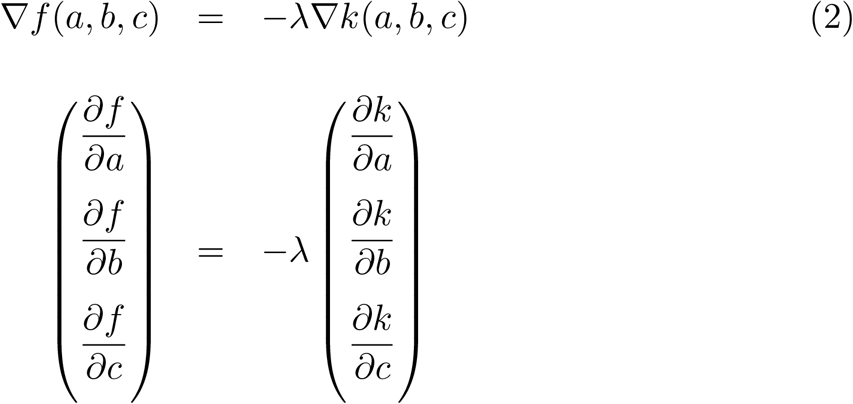

In this case the function *f* (*a, b, c*) = *Total Adhesion Energy* is subject to the condition *k*(*a, b, c*) = *n* = *abc* and *a, b* and *c* are treated as continuous variables.

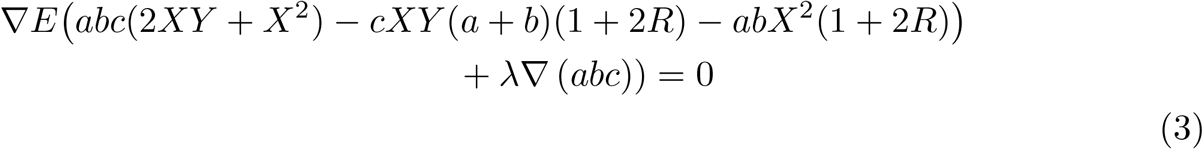

As the relationship holds for each of the vector components, then in conjunction with the condition *k*(*a, b, c*), equation 2 gives a set of simultaneous equations (shown below) whose solutions are critical values (*a*_*c*_, *b*_*c*_, *c*_*c*_), of the constrained function *f* (*a, b, c*).

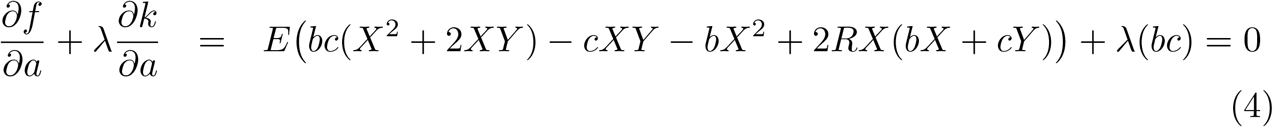

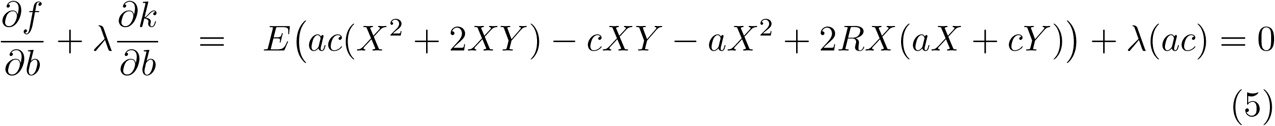

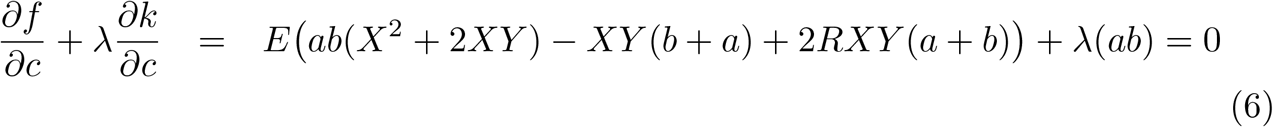

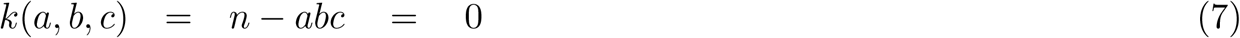

By rearranging equations 4 and 5 for –*λ* and equating them, it can be shown that *a*_*c*_ = *b*_*c*_:

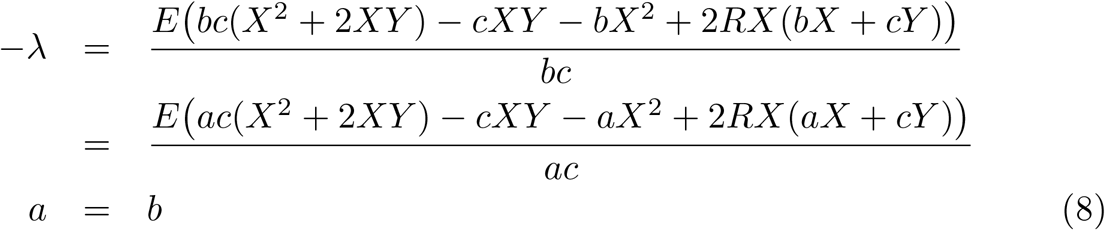

Subscript ‘c’ indicates that the solutions to the simultaneous equations are critical values. Repeating this procedure to eliminate *λ* from equations 4 and 6 provides the information that 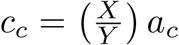:

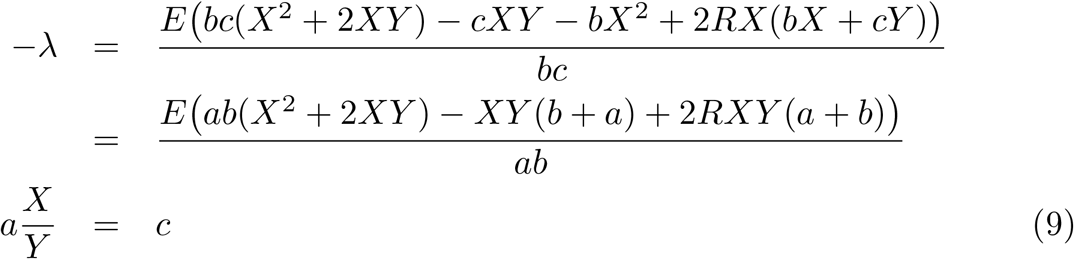

Using the relationships between *a*_*c*_, *b*_*c*_ and *c*_*c*_ and the condition *n* = *abc* it is possible to find an expression for *a*_*c*_, *b*_*c*_ and *c*_*c*_, in terms of *X, Y* and *n*:

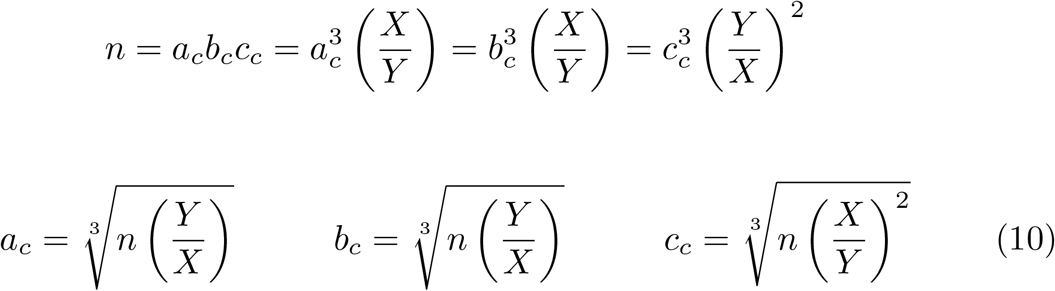

It can be seen that the critical adhesion configuration is not dependent on *R*, the ratio of internal to external adhesion energies per unit area. The information above provides the dimensions of the configuration which is either a maximum, a minimum or saddle point for the total contact area. To determine which type of critical point it is, it is necessary to do a second derivatives analysis. The constrained function for total adhesion energy *g*(*a, b*), for which the critical points have been found, can be expressed explicitly by substituting the expression for *c* from the constraint function into the expression for total adhesion energy *f* (*a, b*):

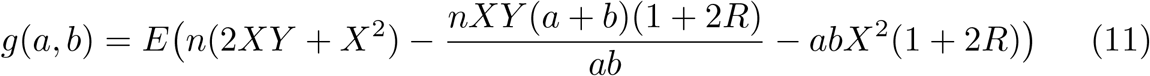

For a function of two variables a partial second derivatives test can determine whether a critical point is maximum, minimum or a saddle point. The conditions for each type of critical point are given below for the constrained total contact area *g*(*a, b*). The conditions above are dependent on the relative rates of changes of second derivatives at the critical points:

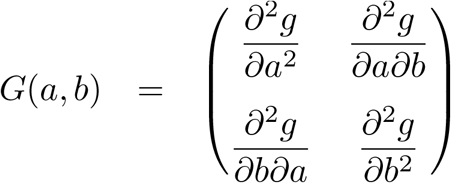

*If det*(*G*(*a*_*c*_, *b*_*c*_)) *>* 0 *and* 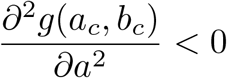 *then* (*a*_*c*_, *b*_*c*_) *is a local maximum*.

*If det*(*G*(*a*_*c*_, *b*_*c*_)) *>* 0 *and* 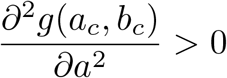 *then* (*a*_*c*_, *b*_*c*_) *is a local minimum*.

*If det*(*G*(*a*_*c*_, *b*_*c*_)) *<* 0 *then* (*a*_*c*_, *b*_*c*_) *is a saddle point*.

*If det*(*G*(*a*_*c*_, *b*_*c*_)) = 0 *then the second derivative test is inconclusive*.

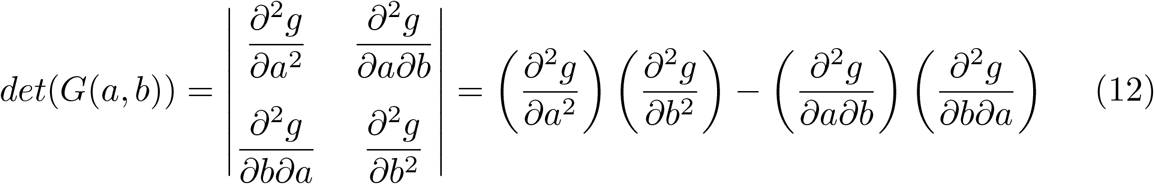

Using the expression for *g*(*a, b*) given in equation 11, the second derivatives can be determined and evaluated at the critical points using expressions for *a*_*c*_ and *b*_*c*_ from equation 10.

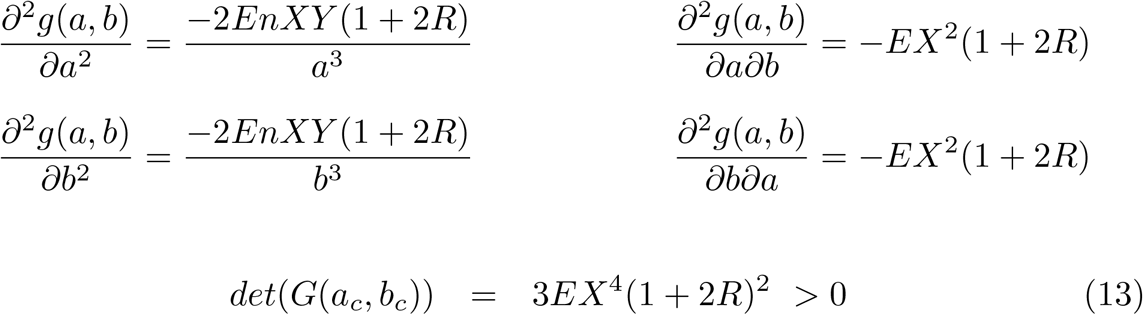

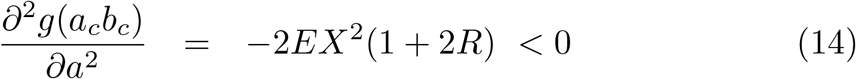

As *E, X* and *R* are real, positive values the critical point (*a*_*c*_, *b*_*c*_, *c*_*c*_) for the constrained total contact area fulfills the criteria for a maximum. Therefore, a population of a finite number of units will have maximum adhesion between the units when the population has a configuration of *a*_*c*_ units by *b*_*c*_ units by *c*_*c*_ units.

It can be seen from equations 10 that the maximum adhesion configuration depends on the ratio of X:Y (width:height) and on the size of the population *n*. When the X:Y ratio is 1:1 (cubic unit structure) the population has a maximum adhesion configuration which is also cubic, with dimensions of 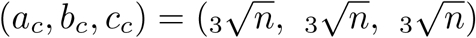 As the unit structure becomes taller and narrower, the maximum adhesion configuration becomes increasingly laminar.

The square-based model described above, and a more general model described in Appendix 2 both assume a discrete number of units lying on a lattice structure. However, the solution for critical values involves treating the population dimensions as continuous variables. This strategy allows for use of the Lagrange multiplier approach to determine constrained maxima, but does not consider edge effects where the configuration of *n* units can not conform to an integer number of units in each dimension. The continuous variable approach provides insight into the overall effect of individual cell adhesive properties on a maximum adhesion-based configuration, but would require refinement to include edge effects if assessment for specific parameter values was necessary.

## Appendix 2

A general model for finding the maximum adhesion configuration of a finite population of cells needs to take into account more parameters than those addressed in the square-based rod model and should also be applicable to different types of cell morphology, cell-cell interactions and cell arrangements. The square-based rod model assumes that adhesion energy is directly proportional to the contact area between two interacting units. However, adhesion is mediated between cells by adhesion molecules which may not be uniformly distributed across the cell, and the morphology of cells is high variable, particularly for neurons. A general model should allow these variations to be taken into account if they are known. For these reasons the general model considers adhesion interaction energies between a population of identical units which are arranged on points of a Bravais lattice structure. A Bravais lattice is a set of points that fills the whole of space and for any choice of position vector **R** the lattice looks the same [3, 4].

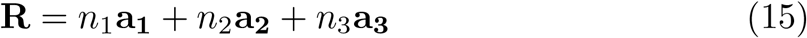

*n*_1_,*n*_2_ and *n*_3_ are integers and **a**_**1**_, **a**_**2**_ and **a**_**3**_ are the primitive vectors of a three dimensional Bravais lattice.

The generalised model has the following further assumptions:

1. Body-, base-, or face-centered lattice structures are excluded and only interactions between points along primitive lattice vectors are considered.
2. The primitive vectors **a**_**1**_, **a**_**2**_ and **a**_**3**_ are of equal length. An adhesion energy is associated with interactions occurring along each of the primitive lattice vector directions. These energies are not length-dependent, and so the lattice structure has the simplest possible length attributes.
3. The configuration of the population can be described as *a* units, by *b* units, by *c* units, along the primitive lattice vector directions. This condition excludes units on a hexagonal lattice structure, where hexagonal configurations need to be considered.
4. Interactions between the population of interest and the external media also occurs along lattice directions, with a reduced adhesion energy compared to internal interactions (ratio *R*¡1).

An example of the general model is shown in figure A1 for a triclinic lattice (**a**_**1**_, **a**_**2**_ and **a**_**3**_ are all non-orthogonal). However, the same model holds for cubic, orthorhombic, and monoclinic lattice structures as there is no dependence on primitive vector lengths or relative angles. There are three types of interaction between neighbors along the axes, with adhesion energies A,B, and C. The adhesion energies can be formulated to include information concerning cell morphology, cell contact areas, distributions of adhesion molecules, and any other factors known to contribute towards adhesion. At this stage only interactions along the axes are included.

**Figure A1:**
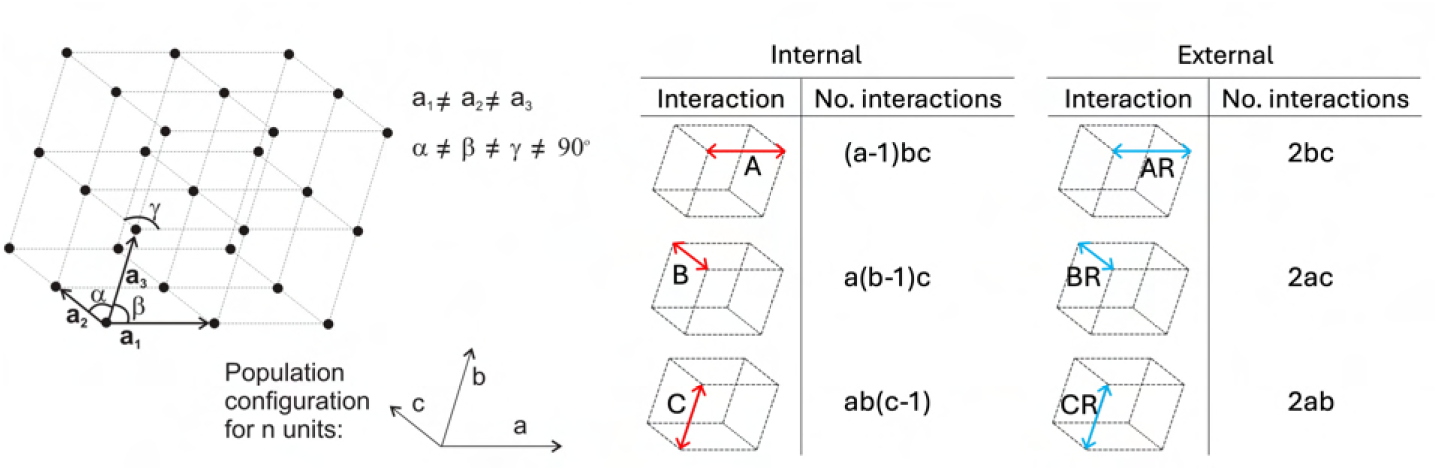
Interactions between units on a triclinic lattice. The numbers of each type of interaction are shown for a population of n units in a configuration with *a* units in the direction of **a**_**1**_, *b* units in the direction of **a**_**2**_ and *c* units in the direction of **a**_**3**_. *A, B* and *C* are adhesion energies associated with interactions that occur along the primitive lattice vector directions (**a**_**1**_, **a**_**2**_ and **a**_**3**_). *R* is the ratio of adhesion energies for external interactions at the edge of the population to internal interactions (within the population)

The configuration of a finite population which maximises the adhesion energy can be found using the same method as for the square-based rods model. The critical points for the total adhesion energy (W) are found using the method of Lagrange multipliers, subject to the constraint that *k*(*a, b, c*) = *n* = *abc*:

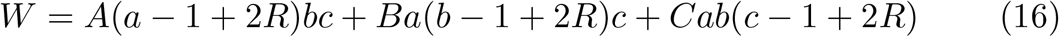

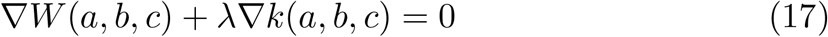

The adhesion energies A,B and C are position invariant as the interaction between two units with the same relative positioning is the same no matter where in the lattice those units are situated. Hence, the adhesion energies are independent of a, b and c. The critical values *a, b* and *c* are given by the solution of the simultaneous equations 18-21 from each component of the Lagrange condition given in equation 17.

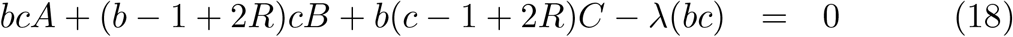

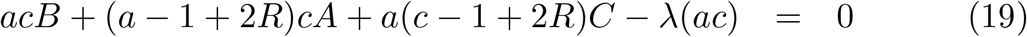

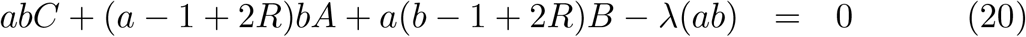

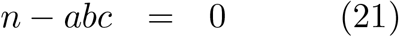

The solution to these simultaneous equations is found in the similar manner to those for the square-based rod model. By eliminating *λ* from equations 18 and 19 and then from 19 and 20, the relationships between *a*_*c*_, *b*_*c*_ and *c*_*c*_ can be determined.

From equations 18 and 19:

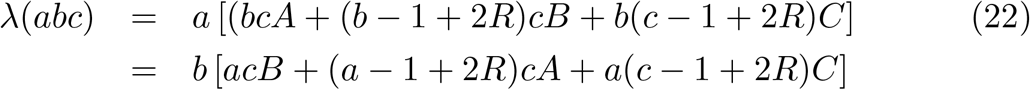

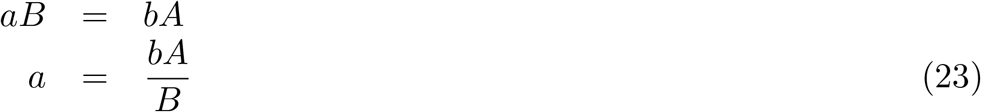

From equations 19 and 20:

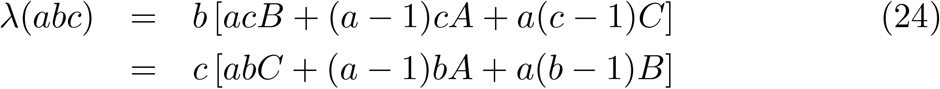

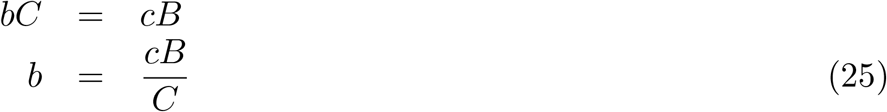

The condition *n* = *abc* reveals the critical points for constrained maximised adhesion:

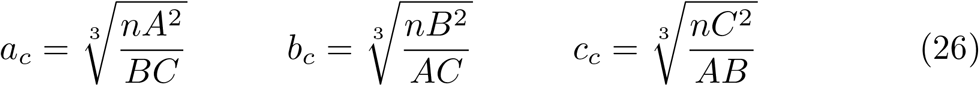

The dimensions for the maximum adhesion configuration is dependent on the ratios of the interaction energies along the three different lattice vectors. The interaction energies A,B and C can be substituted for expressions suitable for a particular situation and the nature of the critical points ascertained using the second order derivatives test. In particular, the general model can be shown to be consistent with the square-based rods model by substituting *A* = *B* = *EXY* and *C* = *EX*^2^, where *E* is the adhesion energy per unit area. Overall, if the adhesion energy increases for interactions given direction relative to the others, then the configuration of the population will increase in the same direction. So for example, if the adhesion energy per interaction increased in the **a**_**1**_ and **a**_**2**_ directions (energies *A* and *B*), relative to energy *C* for interactions in the **a**_**2**_ direction, then the maximal adhesion energy configuration dimension would increase for *a* and *b* and decrease for *c* (more laminar).

